# USP7/Maged1-Mediated H2A Monoubiquitination in the Paraventricular Thalamus: An Epigenetic Mechanism Involved in Cocaine Use Disorder

**DOI:** 10.1101/2022.11.16.516716

**Authors:** Julian Cheron, Leonardo Beccari, Perrine Hagué, Romain Icick, Chloé Despontin, Teresa Carusone, Mathieu Defrance, Sagar Bhogaraju, Elena Martin-Garcia, Roberto Capellan, Rafael Maldonado, Florence Vorspan, Jérôme Bonnefont, Alban de Kerchove d’Exaerde

## Abstract

The individual risk of developing drug addiction is highly determined by the epigenetic landscape^1,2^. Chromatin remodeling regulates drug-induced transcriptional and behavioral effects and the consequent development of addictive behaviors^2,3^. Several chromatin modifications in the ventral tegmental area and nucleus accumbens, including histone H3 methylation, H3 and H4 acetylation, have been implicated in drug addiction. Still, the contribution of other histones and their post-translational modifications (PTMs), such as monoubiquitination is unclear^4–8^. In the course of investigating the underlying mechanisms associated with melanoma-associated antigen D1 (Maged1)^9^, a scaffold protein involved in drug addiction^9^, we found that H2A monoubiquitination in the paraventricular thalamus (PVT) plays a major role in cocaine-adaptive behaviors and cocaine-evoked transcriptional repression. Mice undergoing chronic cocaine administration showed a significant increased monoubiquitination of H2A. Furthermore, we showed that this histone PTM is controlled, in the PVT, by Maged1, along with one of its partner, the deubiquitinase USP7^10^. Accordingly, Maged1 specific inactivation in thalamic vGluT2 neurons, or USP7 inhibition, blocked cocaine-evoked H2A monoubiquitination and abolished cocaine locomotor sensitization. Finally, we identified genetic variations of *MAGED1* and *USP7* associated with modified transition to cocaine addiction and cocaine-induced aggressive behavior in human subjects. These findings identified a new epigenetic modification in a non-canonical reward pathway of the brain and a potent marker of epigenetic risk factor for drug addiction in human.

## Main

Drug addiction, defined as uncontrollable drug intake despite harmful consequences, is a major health problem^11^. The number of deaths associated with cocaine use disorder (CUD) is increasing, and in contrast to tobacco and opioid addiction, no medication has been approved for CUD^11^. Identifying risk factors and therapeutic targets is thus a pressing biomedical challenge. The neurobiological mechanisms that drive and sustain drug addiction involve the combination of neuronal and synaptic plasticity with modifications of gene expression, achieved, in part, through epigenetic mechanisms^1^. By regulating chromatin-related processes, drug-induced epigenetic alterations contribute to the aberrant gene expression and cellular function that underlie drug addiction pathogenesis^2^. Histone post-translational modifications (PTMs) that control chromatin structures regulate drug-adaptive behaviors and the subsequent transition to addiction^3,4^. The promoter of melanoma-associated antigen D1 (*Maged1*) was identified as one of the targets of histone 3 acetylation, a canonical PTM, induced in the (NAc) by chronic cocaine treatment^8^. Subsequently, we identified Maged1 as a master regulator of cocaine reward and reinforcement^9^. Indeed, Maged1 KO mice display, in parallel to an abolished locomotor sensitization, altered cocaine-induced conditioned place preference (CPP) and cocaine self-administration^9^. To date, however, the molecular mechanisms involving Maged1 in these cocaine-related phenotypes have not been characterize^9,12,13^. Despite its ubiquitous expression in the brain and its essential role in cocaine-related behaviors, *Maged1* is not required for cocaine-induced behaviors in dopaminergic or GABAergic cells^9^, leaving excitatory glutamatergic cells as the main candidate.

## Results

### *Maged1* expression in vGluT2 neurons, and specifically in the paraventricular thalamus, is necessary and sufficient for cocaine-related behaviors

Here, we specifically inactivated *Maged1* in glutamatergic vGlut1 or vGluT2 cells by crossing vGluT1- or vGluT2-cre-driver mice with *Maged1* floxed mice (*Maged1^loxP^*). Notably, only vGluT2-mediated (but not vGluT1-mediated) *Maged1* inactivation (*Maged1* conditional knockout (cKO)) abolished cocaine-induced locomotor sensitization (20 mg/kg), a straightforward behavioral paradigm used to model drug-adaptive behavior^14–16^ (Fig. 1a-f and Extended Data Fig. 1). In rodents, repeated cocaine injection induces gradually increased locomotor activity; classically after five days of consecutive injections, the locomotor response reaches a ceiling level. This state lasts for months after cocaine withdrawal^14^. Locomotor sensitization is thus thought to be linked with important aspects of vulnerability to drug addiction^17,18^, relapse and drug craving^19,20^. We further excluded the contribution of glutamatergic telencephalic neurons by using an Emx1-cre driver line (Extended Data Fig. 2). Indeed, specific inactivation of Maged1 in telencephalic Emx1 cells did not alter cocaine-induced locomotor sensitization (Extended Data Fig. 2). Further, to delve into Maged1’s role in the reinforcing properties of cocaine within vGluT2 neurons, we conducted a CPP experiment and operant self-administration assessments (Extended Data Fig. 3). Our CPP results demonstrated a significant distinction between the two groups. Specifically, vGluT2-mediated Maged1 inactivation reduced time spent in the paired compartment associated with cocaine (Extended Data Fig. 3a). This finding suggests that Maged1 have a substantial influence on the response to cocaine-associated stimuli and the development of related adaptive behaviors. Shifting our focus to operant self-administration assessments (Extended Data Fig. 3b-d), we observed that Maged1 inactivation in vGluT2 neurons did not lead to alterations in learning during the training sessions (fixed ratio 1 and 3, Extended Data Fig. 3b-c). However, intriguingly, it did lead to a significant increase in the count of active nose-poking responses during the three consecutive days following learning (refer to supplementary text and Extended Data Fig. 3d). This outcome further supports the notion of Maged1’s potential early involvement in cocaine adaptive behaviors.

**Fig. 1.**
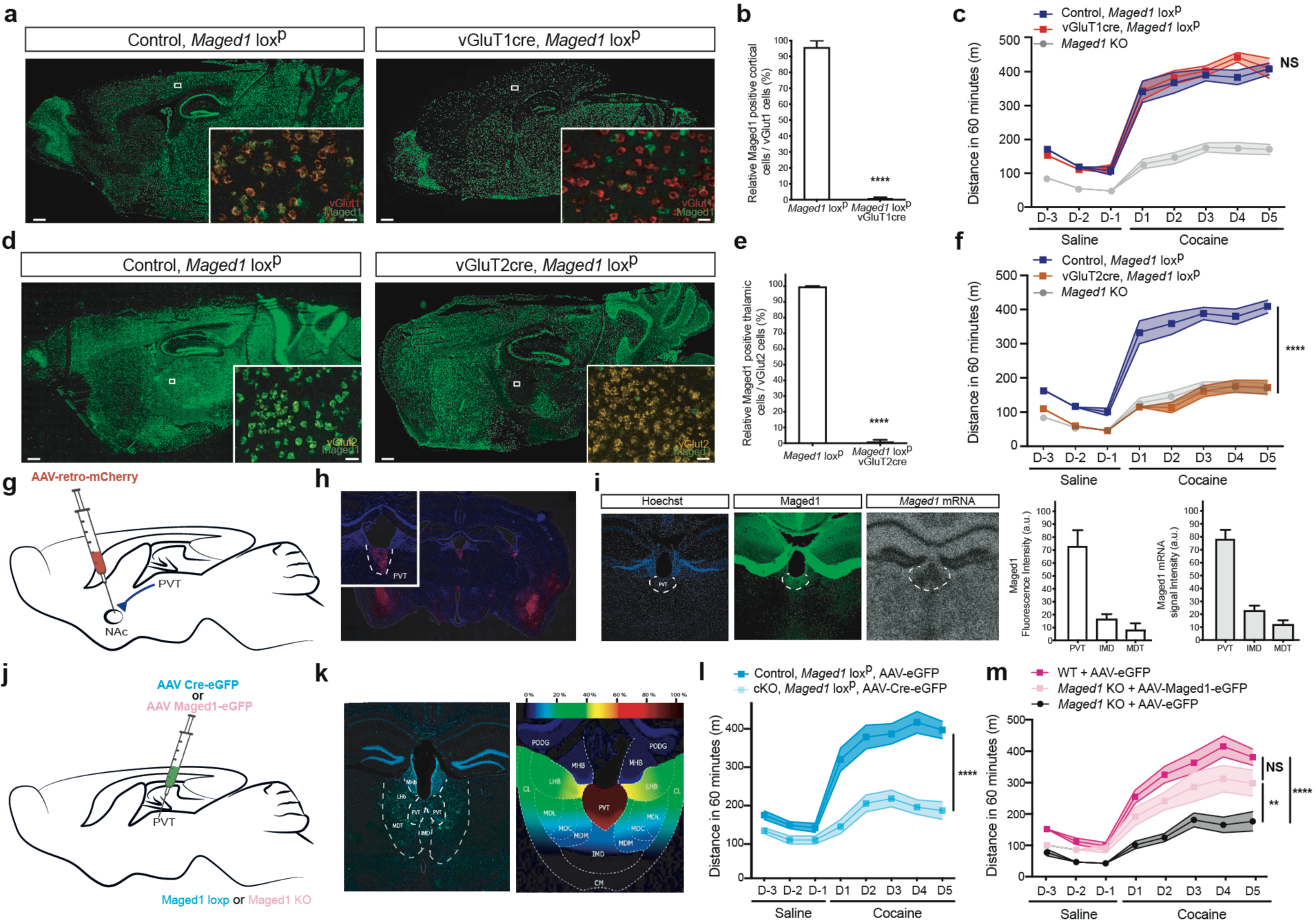
*Maged1* inactivation in vGluT2 neurons completely recapitulates the effect of *Maged1* global inactivation and *Maged1* is ‘necessary and sufficient’ in the PVT. **a,d**, Fluorescent in situ hybridization (FISH) of *Maged1* mRNA (in green) (scale bar: 0.5 mm). Insets show multiplexed FISH with vGluT1 (in red) (**a**), vGluT2 (in orange) (**d**) and *Maged1* mRNA (x20 magnification) in the cortex (**a**) and in the thalamus (**d**) (scale bar: 20 µm (**a**), 30 µm (**d**). **b,e,** Relative percentage of *Maged1* mRNA-positive cells among vGluT1 mRNA-positive cells in the cortex (**b**), and vGluT2 mRNA-positive cells in the thalamus (**e**). n = 7 for each group (Mann– Whitney test, P < 0.0001). **c,f** Cocaine-induced locomotor sensitization, 20 mg/kg, intraperitoneal (ip) injection, Maged1 KO, n = 15. In panel **c**: control, *Maged1* loxP, n = 13; vGluT1cre, *Maged1* loxP mice, n = 7, (two-way ANOVA, control versus vGluT1cre, *Maged1* loxP, P = 0.6651; days, P<0.0001; interaction factor (genotype X days) P = 0.5056). In panel **f**: control, *Maged1* loxP, n = 12; vGluT2cre, *Maged1* loxP mice, n = 9, (two-way ANOVA, control versus vGluT2cre, *Maged1* loxP, P < 0.0001; days, P<0.0001; interaction factor (genotype X days), P < 0.0001). **g,** Schematic of adeno-associated virus (AAV)-retro-mCherry injection in the nucleus accumbens (NAc). **h,** Wide-field image showing the mCherry signal specifically in the PVT and in other NAc-projecting regions, such as the amygdala. Inset shows a clear specific signal in the PVT with a relatively low signal in other thalamic nuclei, demonstrating the relative importance of PVT projecting to the NAc. **i,** Immunohistochemistry (IHC) staining of Maged1 (in green) and Hoechst staining (blue) and parallel in situ hybridization autoradiogram of *Maged1* mRNA and quantification n = 5 mice (for Maged1 fluorescence : Repeated-measure one-way ANOVA, ***p = 0,0002, Tukey’s multiple comparisons test, PVT vs IMD, **P = 0.0027; PVT vs MDT, **P = 0.002; IMD vs MDT P = 0.1552; for Maged1 mRNA : Repeated-measure one-way ANOVA, ****p < 0.0001, Tukey’s multiple comparisons test, PVT vs IMD, ***P = 0.0007; PVT vs MDT, ***P = 0.0002; IMD vs MDT P = 0.0734). **j,** Schematic of inactivation (AAV-Cre-eGFP into *Maged1* loxP mice) and re-expression experiments in the PVT (AAV-*Maged1*-eGFP in the *Maged1*-KO). **k,** IHC staining showing the eGFP signal (in green) in the PVT (-2 mm from the bregma) (left), topographic representation of the targeted area; colors represent the percentage of the superimposed eGFP-targeted area (right). **l,m,** Inactivation experiment (**l**) and re-expression experiment (**m**) constisting of a cocaine-induced locomotor sensitization, 20 mg/kg, ip injection. In panel **m**: control, *Maged1* loxP, AAV-eGFP mice, n = 10; cKO, *Maged1* loxP, AAV-Cre-eGFP mice, n = 11, (two-way ANOVA, control versus *Maged1*-cKO, P < 0.0001; days, P<0.0001; interaction factor (genotype X days) P < 0.0001). In panel **m**: control *Maged1*-KO, AAV-eGFP mice, n = 10; control *Maged1* WT, AAV-eGFP mice, n = 10; re-expressed in KO, AAV-*Maged1*-eGFP mice, n = 8, (two-way ANOVA, groups P < 0.0001; days, P<0.0001; interaction factor (genotype X days) P < 0.0001 and Sidak’s post-test; re-expressed in KO versus control *Maged1* WT mice, P = 0.0731; re-expressed in KO versus control Maged1-KO mice, P = 0.0072; control *Maged1* WT versus control *Maged1*-KO, P < 0.0001). All error bars represent s.e.m. PVT: the paraventricular nucleus of the thalamus, MDT: the mediodorsal thalamic nucleus (including the medial, lateral and central parts), IMD: the intermediodorsal nucleus of thalamus.

To screen for potential vGluT2 nuclei involved in acquisition of this phenotype, we performed AAV retro mCherry injection in the NAc and revealed the amygdala, the ventral subiculum and the paraventricular thalamus (PVT) as the main vGluT2 nuclei targeting the NAc (Fig. 1g,h). Of these, *Maged1* mRNA in situ hybridization and immunohistochemistry (IHC) showed a singularly high level of Maged1 expression (Fig. 1i) in the PVT, which has recently been shown to be involved in drug addiction^21–24^. We therefore examined the impact of *Maged1* inactivation in the PVT via stereotactic injection of an adeno-associated virus (AAV) encoding a Cre recombinase in adult *Maged1^loxP^* mice (Fig. 1j,k). Here again, we observed the abolition of cocaine-induced locomotor sensitization; thus, phenocopying *Maged1*-KO mice and excluding developmental effects (Fig. 1l). To test the sufficiency of Maged1 in the PVT to cocaine sensitization, we stereotactically injected an AAV expressing Maged1 into *Maged1*-KO mice and found that cocaine-evoked sensitization was restored (Fig. 1j,k,m). In contrast, stereotactic inactivation^9^ (Extended Data Fig. 4) or re-expression (Extended Data Fig. 5) of *Maged1* in other vGluT2- or mixed vGluT1/vGLuT2 cell hubs of drug addiction, such as the prefrontal cortex (PFC), amygdala or ventral subiculum^25^, did not lead to major significant changes in cocaine-induced locomotor sensitization. Taken together, these results point toward PVT as the key nucleus where Maged1 plays a role in drug addiction.

### Cocaine-induced transcriptomic changes imply *Maged1*-mediated Polycomb repressive complex

To identify the molecular mechanisms linking Maged1 in the PVT with cocaine-mediated behavior, we analyzed the transcriptome in the thalamus after chronic cocaine (or saline) administration in control and *Maged1*-cKO mice by RNA sequencing (RNA-seq) (Fig. 2a). In control mice, repeated cocaine injection induced upregulated expression of 252 genes and downregulated expression of 452 genes compared to the expression of these genes after saline treatment (Fig. 2b). The transcriptional response (and particularly the downregulation process, see supplementary text) to cocaine exposure by *Maged1*-cKO mice differed from that of control mice in terms of genes and their assigned GO terms (Fig. 2b and Extended Data Fig. 6 and 7a, and Table 1). Interestingly, a gene set enrichment analysis with curated gene sets revealed that cocaine-regulated genes in control but not in *Maged1*-cKO mice were targets of the Polycomb repressive complex (PRC) (Fig. 2c,d and Extended Data Table 1). Briefly, PRC2 mono-, di-, and tri-methylates lysine 27 of histone H3 (H3K27me1/me2/me3)^26,27^, and PRC1 monoubiquitinates histone H2A on lysine 119 (H2AK119ub1)^28^ (Fig. 2d). Notably, the genes with expression similarly regulated by cocaine in the control and *Maged1*-cKO mice were not known to be targeted by PRC action, as we did not find any significant overlap between genes known to be targeted by PRC and these Maged1-independent genes.

**Fig. 2.**
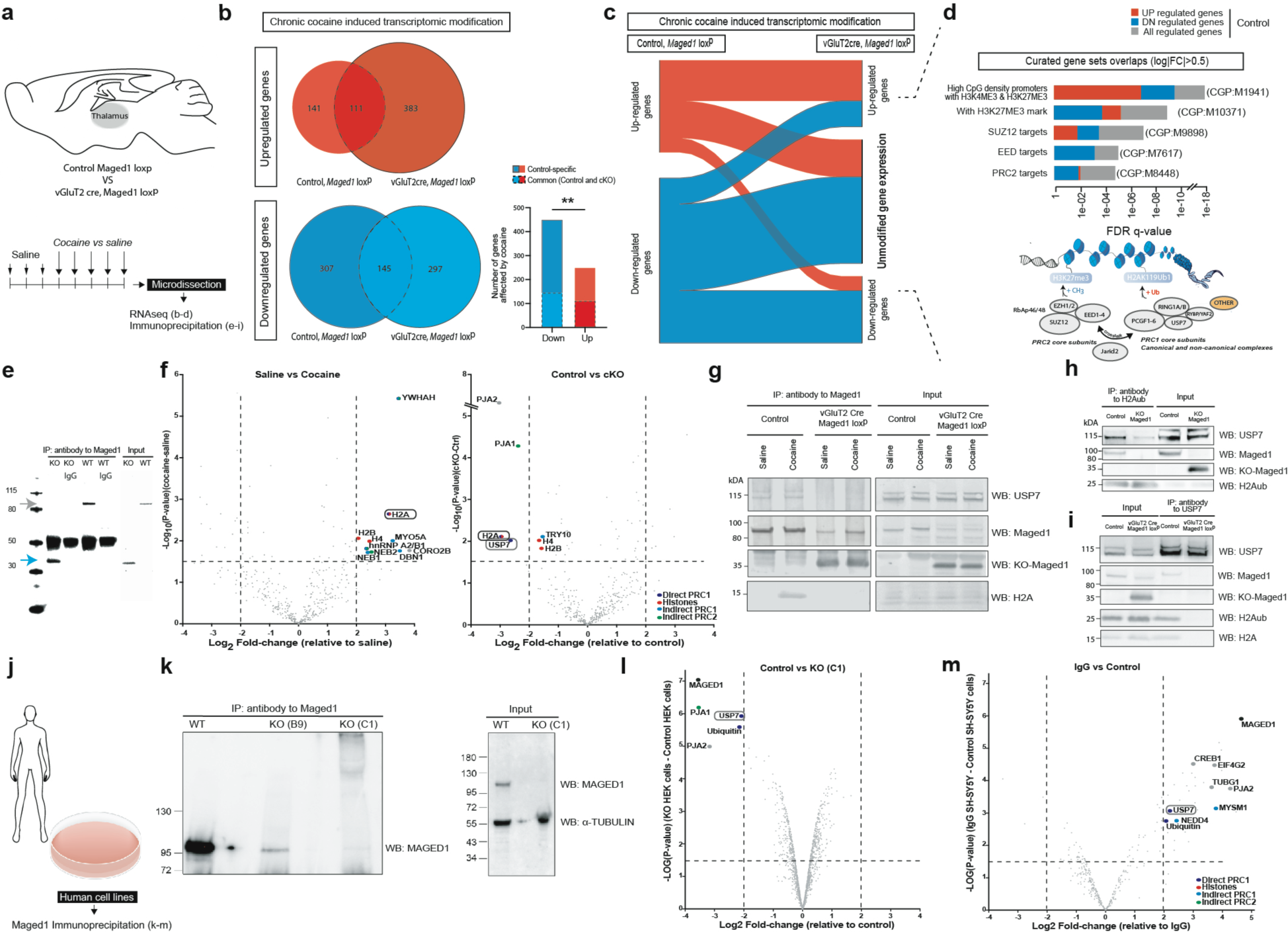
*Maged1* inactivation modifies gene expression regulated by the Polycomb pathway through the USP7-H2A interaction. **a,** Method and time line for RNA sequencing and immunoprecipitation (IP) from microdissected mouse thalami. **b**, Venn diagram showing significantly up- and downregulated genes in control mice, *Maged1* loxP mice and *Maged1*-cKO, vGluT2 cre, *Maged1* loxP mice (log2|FC|≥0.5, FDR<0.05). The histogram on the bottom right shows that the proportions of overlapping up- and downregulated genes between control and *Maged1*-cKO group differ (chi-square, **P < 0.01). **c,** Alluvial plot showing the proportion of genes that are up- or downregulated in the control, *Maged1* loxP and are or are not modified in *Maged1*-cKO mice, vGluT2 cre, *Maged1* loxP mice (log2|FC|≥0.5, FDR<0.05). **d,** Curated gene set analysis showing statistically significant overlapping with literature-based gene categories and the unmodified genes in *Maged1*-cKO mice (n = 448 genes changed in the control) (top) and scheme of the Polycomb repressive complexes, PRC1 and PRC2, and their crosstalk (bottom). **e,** Control antibody (IgG) or anti-Maged1 antibody-based (IP) with in vivo thalamic tissue; anti-Maged1 antibody was used for western blotting (WB) Maged1 band is indicated with a gray arrow, Maged1-KO truncated protein is indicated with a blue arrow. **f,** Mass spectrometry (MS) results, for every identified protein, log2 FC values are plotted on the X-axis, and -log10 p values are plotted on the Y-axis of the volcano plot. On the left, relative abundance of proteins between Maged1 IP from control mice treated with saline (n = 5) and control mice treated with cocaine (n = 7), and on the right, between control mice (n = 7) and *Maged1*-cKO, vGluT2cre, *Maged1* loxp (n = 8) mice (both treated with cocaine) is shown. **g,** Mononucleosome (Mn) IP of Maged1 and its partners with USP7 and H2A identification by WB. **h,** MnIP of monoubiquitinated H2AK119 (H2Aub) and its partners with identification of Maged1 and USP7 by WB. **i,** MnIP of USP7 and its partners with identification of Maged1 and H2Aub and H2A by WB. Panels **h** and **i** were performed under saline treatment conditions but are also representative of MnIP under cocaine treatment conditions. **j,** Method for IP with human cells (panels **k,l,m**). **k,** western blot (WB) analysis of wild type (WT) and C1 subclones of HEK293T lysate immunoblotted with anti-MAGED1 and anti-α-TUBULIN antibodies (left), WB showing IP of endogenous MAGED1 in WT, B9 and C1 subclones of HEK293T lysates with an anti-MAGED1 antibody (right). **l,m** MS results represented by volcano plots showing the relative abundance of proteins between MAGED1 IP from WT versus (C1)-KO HEK293T cells (**l**) and in the WT versus rabbit IgG (isotype control) in SH-SY5Y cells (**m**). PRAJA1/2, CREB1, NEDD4 (E3 ubiquitin ligase) and the PRC1 deubiquitinase USP7 are the main interactors of MAGED1. For MS results (**f,l,m**), proteins above the specified threshold (-log10 p value > 1.5 and log2|FC| > 2) are highlighted with their names and a color code (a direct PRC1 component is dark blue, histones are red, known PRC1 interactors are blue and known PRC2 interactors are green). Histone H4, H2B and Try10 are highlighted because of their proximity to the thresholds and their links with the H2A-PRC1 pathway.

### Maged1 enables interaction between histone H2A and USP7

To understand how Maged1 mediates the profound transcriptional changes observed upon cocaine exposure (Fig. 2b-d), we aimed at identifying Maged1 protein partners in the thalamus by performing coimmunoprecipitation and mass spectrometry (co-IP-MS) experiments with an anti-Maged1 antibody that recognizes both the full length and truncated form (functional KO) in saline- and cocaine-treated mice (Fig 2e). We identified 404 potential Maged1-interacting proteins (see Supplementary Text and Table 2). Among them, histone H2A, in its abundantly expressed H2A.2 variant in the adult brain^29,30^, and de-ubiquitinase USP7, a well-characterized PRC1 component^10,31,32^, as well as several other proteins indirectly linked to PRC complexes, such as Praja1, hnRNPA2/B1, Myo5a, Dbn1 and Neb1/2, were identified (Fig. 2f)^33,34^. Within all the identified candidates, histone H2A was the only one whose i) interaction with Maged1 was disrupted in the *Maged1*-cKO samples and ii) displayed a differential association with Maged1 in cocaine and saline treated control mice (Fig 2f and Extended Data Table 2). USP7 behaved as a Maged1 partner under both saline and cocaine treatment conditions (Fig. 2f and Extended Data Table 2). To confirm these interactions, we performed Maged1 co-IP with mononucleosome samples extracted from the thalamus, followed by western blotting (WB) for protein identification. We confirmed that the specific link between Maged1 and H2A was evident only in the presence of cocaine, while USP7 coprecipitated with Maged1 under both cocaine and saline treatment conditions (Fig. 2g). In agreement with these IP data, we observed that Maged1 and USP7 protein levels were both higher in the PVT than in other parts of the thalamus, like the adjacent mediodorsal thalamic nucleus (MDT) (including its medial, lateral and central parts) and the intermediodorsal nucleus of thalamus (IMD). (Extended Data Fig. 8).

Next, we postulated that the Maged1 interaction with H2A and USP7 contributes to the regulation of the monoubiquitination of H2A at lysine 119 (H2AK119ub1 (hereafter H2Aub)) as Mage proteins assemble with different E2/3 ubiquitin ligases^13,35^. This histone mark deposited by PRC1 and linked to gene repression^36,37^ may explain how Maged1 represses gene expression in thalamic neurons upon chronic cocaine administration. To test this hypothesis, we performed co-IP experiments using an anti-H2Aub monoclonal antibody and observed that USP7 and Maged1 both linked with H2Aub under saline and cocaine treatment conditions (Fig. 2h and Extended Data Fig. 9). Notably, USP7 binding to H2Aub was significantly decreased in the *Maged1*-KO mice, indicating that Maged1 was necessary for this H2Aub-USP7 interaction (Fig. 2h and Extended Data Fig. 9). Furthermore, USP7-specific antibody coimmunoprecipitated H2Aub and H2A under saline and cocaine treatment conditions but not in the Maged1-cKO thalamus samples (Fig. 2i). From these newly demonstrated interactions we hypothesized that Maged1 plays a role in H2A monoubiquitination in thalamic neurons following cocaine intake, via Maged1 involvement with a PRC1 member. To test whether MAGED1 associates with USP7 in human cells, we performed co-IP-MS of MAGED1 in two different human cell lines: human embryonic kidney 293T (HEK293T) and neuroblastoma (SH-SY5Y) cells (Fig. 2j-m and Extended Data Table 2). The interaction with USP7 was thus confirmed in human cells (Fig. 2l,m). In neuronal cells, the known interaction between Maged1 and CREB^38^ was also found (Fig. 2m). In addition, a novel interaction between MAGED1 and another H2A deubiquitinase, MYSM1, was also identified (Fig. 2m). These results indicated that the Maged1-USP7 axis is evolutionarily conserved and may regulate addiction behavior in humans through histone H2A PTM.

### Cocaine induces an increase of histone H2A monoubiquitination in the NAc, PFC and thalamus

We therefore tested whether chronic administration of cocaine impacts H2A monoubiquitination in key brain regions involved in drug addiction and whether these changes depend on Maged1 function (Fig. 3a).To this end, by WB using anti-H2Aub and anti-H2A specific antibodies, we analyzed H2A and H2Aub levels in the PFC, thalamus and NAc of control and *Maged1*-cKO animals after chronic cocaine or saline administration. The results showed a significant increase in H2Aub in all these nuclei (Fig. 3b-e) after cocaine administration. Remarkably, the cocaine-dependent increase in H2Aub was only abolished in the thalamus of the *Maged1*-cKO mice (Fig. 3e). These results are the first to show that cocaine treatment induces an increase in H2Aub in key structures involved in addiction and that Maged1 is crucial for cocaine-induced H2A monoubiquitination, specifically in the thalamus.

**Fig. 3.**
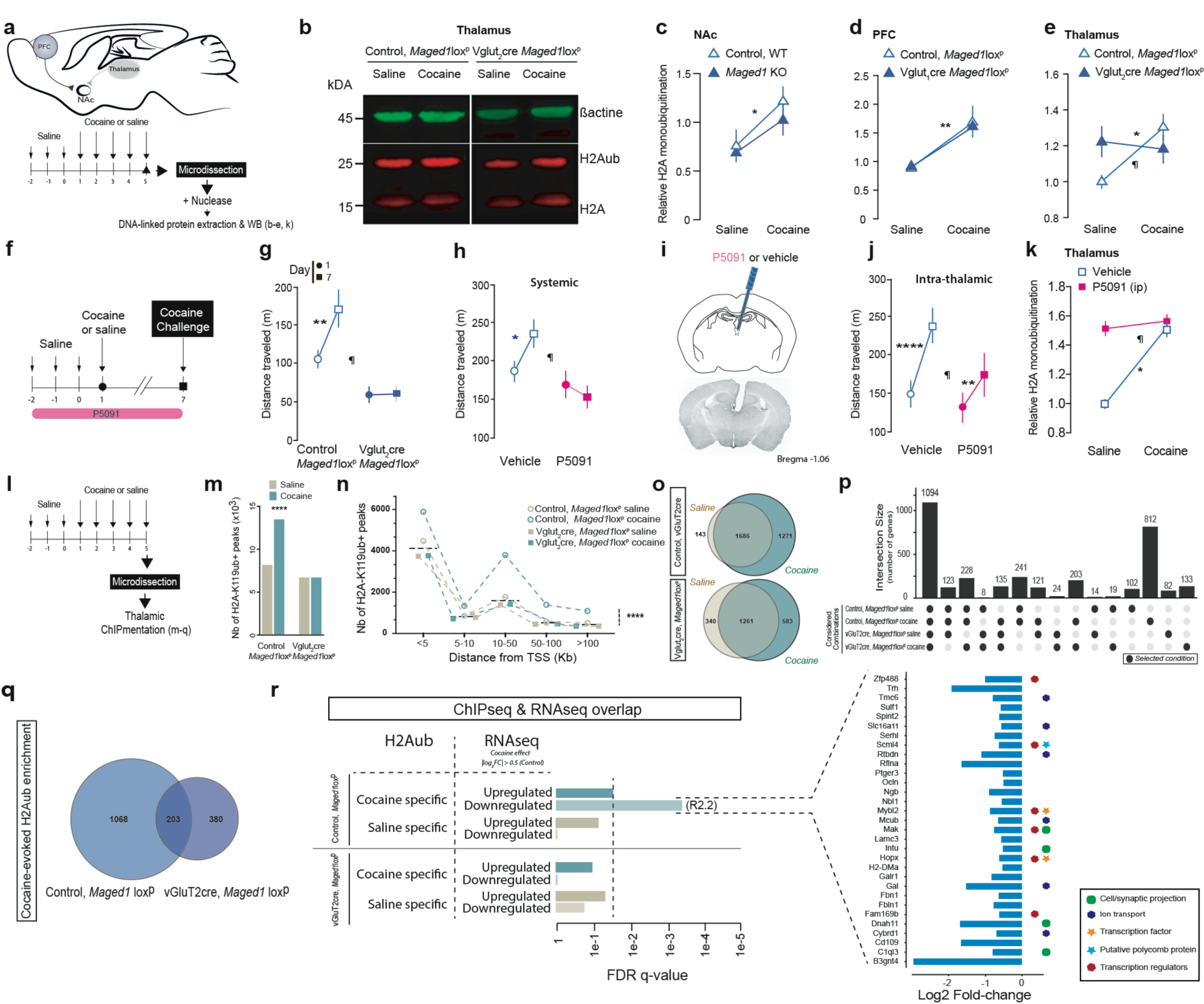
Chronic cocaine administration increases H2A ubiquitination, which is impaired by *Maged1* inactivation or UPS7 inhibition. **a,** Method and time line of the experiment (cocaine, intraperitoneal (ip) injection, 15 mg/kg). Protein extraction was performed with nuclease to release protein linked to DNA. **b,** Representative fluorescence western blot (experiments performed with thalamus samples). **c,** Relative H2A monoubiquitination (H2AK119ub1, and 2Aub) in the NAc (**c**), in the PFC (**d**) and in the thalamus (**e**), quantified as the ratio between H2Aub and H2A signal intensities after saline or cocaine administration. In panel **c**: control saline, n = 6; control cocaine, n = 5; cKO saline, n = 5; cKO cocaine, n = 4, (two-way ANOVA, control versus *Maged1*-cKO, P = 0.3839; saline versus cocaine, *P = 0.0144; interaction factor, P = 0. 6853). In panel **d**: control saline, n = 9; control cocaine, n = 11; cKO saline, n = 10; cKO cocaine, n = 8, (two-way ANOVA, control versus *Maged1*-cKO, P = 0.8751, saline versus cocaine, **P = 0.0023; interaction factor, P = 0. 8428). In panel **e**: control saline, n = 14; control cocaine, n = 20; cKO saline, n = 13; cKO cocaine, n = 1, (two-way ANOVA, control versus *Maged1*-cKO, P = 0.0849; saline versus cocaine, P = 0.4907; interaction factor, ¶P = 0.0246, Sidak’s post-test (saline versus cocaine), *P = 0.0281 for the control group). **f,** Scheme of cocaine sensitization (cocaine, ip injection, 15 mg/kg) during the USP7 inhibition experiment (P5091, ip injection, 10 mg/kg; intrathalamic injection, 0.1 mg/kg, and 400 nl). **g,** Cocaine-induced sensitization was abolished after inactivation of *Maged1* in vGluT2 neurons (*Maged1*-cKO, vGluT2cre *Maged1* loxp), control, n = 9; cKO, n = 8, (two-way ANOVA, control versus cKO, P = 0.0007; days, P < 0.0216; interaction factor (genotype X days) ¶P = 0.0249; Sidak’s post-test (D1 versus D7) **P = 0.0044). **h,** Cocaine-induced sensitization was abolished after systemic inhibition of USP7, control, n = 10; P5091, n = 8; (two-way ANOVA, vehicle versus P5091, P = 0.0224; days, P = 0.1790; interaction factor (P5091 X days) ¶P = 0.0125; Sidak’s post-test (D1 versus D7) * P = 0.012). **i,** Diagram showing cannula implantation for injecting P5091 into the paraventricular thalamus (PVT) and immunohistochemical staining for Maged1. **j,** Cocaine-induced sensitization was significantly diminished after intra-PVT P5091 injections, control, n = 7; P5091, n = 8, (two-way ANOVA, vehicle versus P5091, P = 0.1987; days, P < 0.0001; interaction factor (P5091 X days) ¶P = 0.0265; Sidak’s post-test (D1 versus D7), ****P < 0.0001, ** P = 0.011). **k,** Relative H2Aub in the thalamus, control saline, n = 10; control cocaine, n = 4; P5091 saline, n = 6; P5091 cocaine, n = 9 (two-way ANOVA, vehicle versus P5091, P = 0.0007; saline versus cocaine, P = 0.0171; interaction factor ¶P = 0.0414; Sidak’s post-test (saline versus cocaine) *P = 0.0249 for control group). **h,j,k**, All control mice received vehicle injections. The distance traveled during cocaine-induced sensitization was recorded for 30 minutes. **l,** Time line of the experiment before microdissection of the thalamus used for ChIPmentation. **m,n,** Plots representing the number of H2Aub+ elements identified across different experimental conditions (**m**) and their distribution with respect to the nearest gene transcription start site (TSS) (**n**). The blue dashed line in panel **m** represents the median of 4 different experimental conditions. Repeated cocaine injection was associated with an increase in the total number of H2Aub+ elements compared to the saline condition; this increase was not observed in *Maged1*-cKO mice (chi-square, ****P<0.0001). **o-q**, Euler plots (**o,q**) and Upset (**p**) representing the number of H2Aub+ genes (displaying H2Aub+ elements at their promoters) across the different experimental conditions. Cocaine injection in control mice resulted in an increase in H2Aub+ genes (**o**, top). In contrast, in *Maged1*-cKO mice, the number of cocaine-specific H2Aub + genes was considerably lower than that in control mice (**o**, bottom), and only a minor fraction of cocaine-specific H2Aub+ genes were shared between control and *Maged1*-cKO mice (**q**). The Upset plot (**p**) represents the number of H2Aub+ genes in common between the different experimental groups; bars represent the number of H2Aub+ genes in common (Intersection size) between the groups indicated below each of them with a dark grey circle. **r,** Bar plot presenting the overlapping genes displaying H2Aub enrichment in saline or cocaine condition (in *Maged1* control and cKO mice) and up- or downregulated expression in *Maged1* control mice (log2|fold change, FC|≥0.5, false discovery rate (FDR)<0.05). A significant overlap was found between genes displaying H2Aub enrichment and that are downregulated after cocaine administration. These genes are highlighted on a bar plot with their associated log2FC in control mice on the right.

### USP7 pharmacological inhibition or Maged1 inactivation precludes cocaine-evoked H2A monoubiquitination and cocaine sensitization

Because USP7 is a known PRC1 regulator with deubiquitinase activity^10,31,32^, we hypothesized that USP7 plays a key role in the Maged1-dependent cocaine behavioral response. We therefore inhibited USP7 activity in vivo by administrating P5091, a selective USP7 inhibitor, in a two-injection cocaine sensitization protocol^15^ (Fig. 3f). Interestingly, systemic injection of P5091 abolished locomotor sensitization in control mice, similar to the effect of *Maged1* inactivation (Fig. 3g,h and Extended Data Fig. 10), with no effect on spontaneous locomotor activity (Extended Data Fig. 10b), and no effect on the locomotor activity of Maged1 KO mice (Extended Data Fig. 10c). To exclude a transient occlusion effect, we continued the experiment for an additional 3 days of daily cocaine injections and observed a sustained abolition of cocaine sensitization (Extended Data Fig. 10d). Similarly, an intrathalamic injection of P5091 significantly interfered with cocaine-induced locomotor sensitization (Fig. 3i,j). Similar to the observation with *Maged1*-cKO mice, P5091 treatment prevented the increase in H2A monoubiquitination induced by cocaine administration (Fig. 3k). These results showed that pharmacological abolition of cocaine-evoked H2Aub deposition in the thalamus paralleled an alteration of cocaine-evoked behavioral sensitization.

### Cocaine-evoked histone H2A monoubiquitination and its subsequent transcriptional repression

To further characterize the mechanisms by which Maged1 activity and H2A mono-ubiquitination contribute to cocaine sensitization, we compared H2Aub genomic coverage in the thalamus of control and *Maged1*-cKO mice after chronic cocaine or saline were administered by performing chromatin immunoprecipitation with sequencing (ChIPmentation)^39^ (Fig. 3l). Analysis of the H2Aub coverage in these mice revealed a higher number of regions with this PTM in the cocaine-treated mice compared to saline-treated mice (13493 and 8174, respectively; +164.95% in the cocaine vs. the saline treated mice) (Fig. 3m). Supporting a regulatory function of this PTM, the same increase was observed when only gene promoter regions (defined as 5kb upstream and 2k downstream of the gene transcription start site (TSS)) were analyzed, and the largest fraction of H2Aub+ elements was located less than 5 kb upstream from the nearest TSS. Cocaine shifted this distribution significantly upward in the control but not in the *Maged1*-cKO mice (Fig 3n). Specifically, 1271 genes with H2Aub at their promoters were found in the control mice in the cocaine treatment condition, while only 143 loci showed H2Aub specific marks after saline administration (Fig. 3o). In contrast, the total number of H2Aub marks and promoters was comparable between the cocaine-injected (583 genes) and saline-injected (340 genes) *Maged1*-cKO animals (Fig. 3o). Furthermore, 84.03% (1068 genes) of genes displaying H2Aub enrichment upon cocaine administration in control mice did not show enrichment in *Maged1*-cKO animals, indicating that the loss of *Maged1* deeply impacted the H2Aub dynamics associated with the cocaine response (Fig. 3p,q). Cocaine-dependent H2Aub+ genes in control mice were associated with changes in biological process like ion transport, synaptic signaling, axonal guidance, neuron projection and cytoskeleton/microtubule organization (Extended Data Fig. 11 and Table 1) which were not found in *Maged1*-cKO mice (Extended Data Table 1). These results support the idea that Maged1 exerts its specific effects on the behavioral response to cocaine primarily through drug-induced PRC1 gene repression, as cocaine administration to control mice triggered coherent transcriptional and H2Aub dynamics (Extended Data Fig. 6d,7,11). Indeed, we found a significant overlap, between RNAseq and ChIP data, in the genes with transcriptional downregulation and those with promoters enriched in H2Aub marks after cocaine administration (Fig. 3r). Interestingly, we found that c1qL3, a signaling protein involved in synaptic projection and human CUD^40,41^ was downregulated while displaying an increase in H2Aub on its promoter upon chronic cocaine administration, as well as HOPX and SCML4, important epigenetic regulators linked to PRC1.

### MAGED1-USP7 modifies addiction risk and behavioral response to cocaine in human

To assess the potential contributions of *MAGED1* and *USP7* to CUD directly in humans, we conducted a within-case *MAGED1* and *USP7* gene analysis with 351 consecutively recruited outpatients with cocaine addiction and genetically verified Caucasian ancestry. After demonstrating the absence of structural variants in/near *MAGED1* or *USP7*, we focused on 175 nucleotide polymorphisms (SNPs). We investigated their potential associations with seven CUD-related phenotypes reflecting all stages of disease progression, correcting for multiple testing. The outpatient population comprised 77% men aged 38 +/- 9 years old. All the outpatients reported lifetime cocaine use and 72% of these men suffered from severe CUD (including to crack-cocaine) (Extended Data Table 3). Comorbid substance use disorders (SUDs) were tobacco smoking (current for 89% of the sample), alcohol (62%), opioid (61%), cannabis (64%), and/or benzodiazepine (41%) use disorders (Extended Data Table 3). Half the sample had three lifetime SUDs or more (tobacco excluded). Patients carrying at least one alternate allele were compared to those carrying only reference allele(s). In the case of multiple statistical associations with a given phenotype, we retained lead SNPs most likely associated with the underlying biological signal by combining positional and functional data. First, the transition from 1^st^ cocaine use to CUD was significantly associated with 17 SNPs in *MAGED1* (Fig. 4a,b), of which we identified two lead SNPs. These two SNPs showed opposite effects during the transition, as highlighted by Kaplan–Meier survival curves (Fig. 4c). An *in silico* analysis with human cells showed a significant association of these two SNPs with DNA methylation changes and thus altered gene expression. Second, we observed associations between 44 *USP7* SNPs and cocaine-induced aggression (Fig. 4a), of which we identified five lead SNPs. Four SNPs were in the alternate allele associated with lower aggression scores, and scores were higher for the alternate allele of the other SNP (Fig. 4c). Consistent with our preclinical data, these findings with human samples revealed that *MAGED1* and *USP7* polymorphisms are associated with CUD (Extended Data Table 4). These associations were independent of potential confounders, such as the self-reported amount of cocaine use, which is strongly related to cocaine-induced aggression and childhood hyperactivity and is associated with a shorter transition from first cocaine use to CUD (Extended Data Table 5 and 6). These data enabled us to designate *MAGED1* and *USP7* as contributors to the vulnerability for CUD and specific behavioral responses to cocaine.

**Fig. 4.**
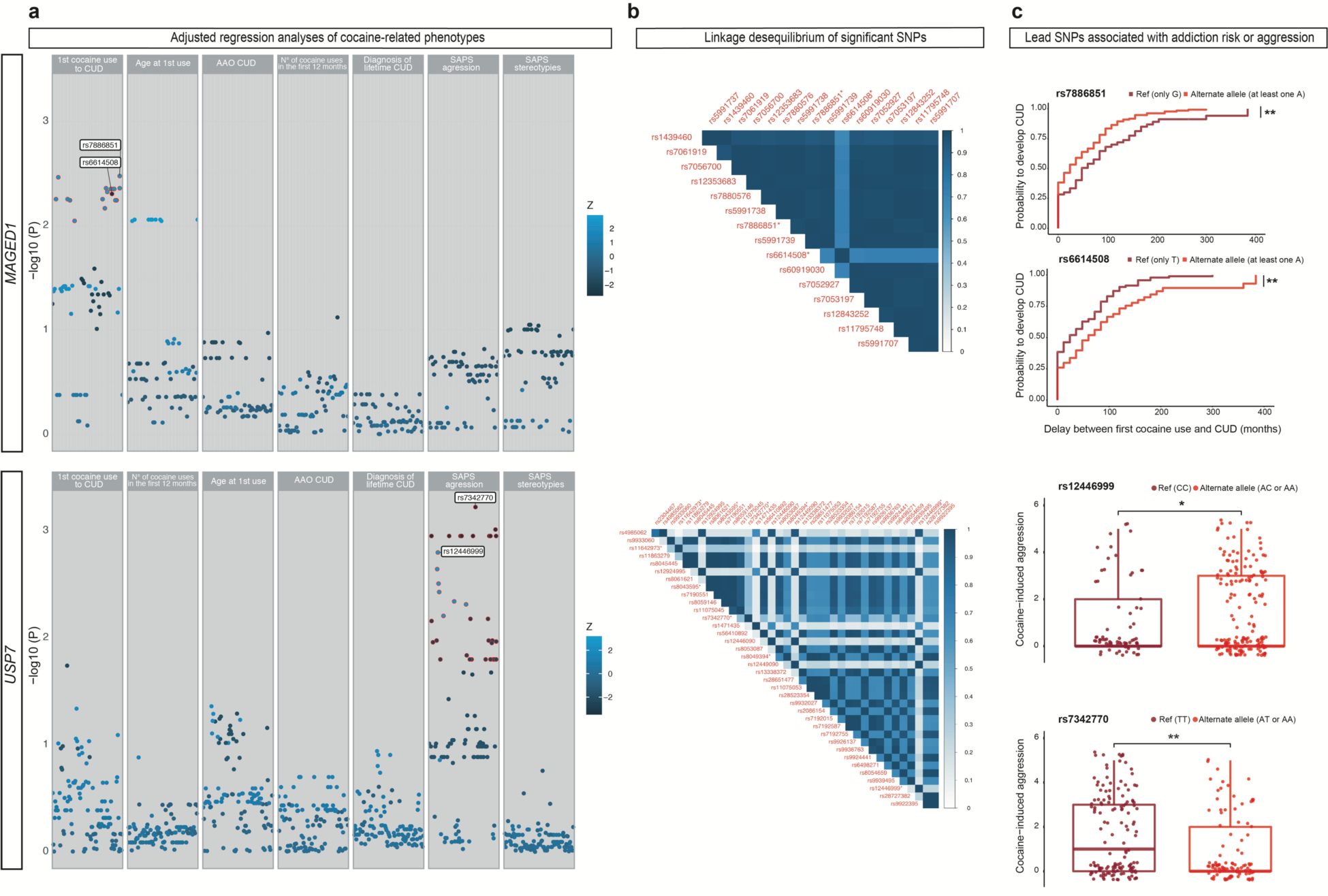
Cocaine use disorder (CUD) is associated with small *MAGED1* and *USP7* polymorphisms in polydrug users. **a,** Scatterplots representing -log (p values) on the y-axis and effect sizes (Z) as the point color for 61 *MAGED1* single nucleotide polymorphisms (SNPs, upper panel) and 114 *USP7* SNPs (lower panel) tested using regression analyses with seven cocaine-related phenotypes (Cox regression for age at 1^st^ cocaine use and transition from 1^st^ cocaine use to cocaine user disorder (CUD), linear for the other), adjusted for sex and ancestry of 351 patients with strict Caucasian ancestry. Significant SNPs are circled in red, and the two most significant in each phenotype are labeled with their name. **b,** Linkage disequilibrium heatmaps showing increasing r^2^ values, as indicated from white to dark blue with 16 *MAGED1* SNPs (data were not available for one SNP) significantly associated with the transition from 1^st^ cocaine use to CUD (upper panel) and 37 *USP7* SNPs (data were not available for seven SNPs) significantly associated with cocaine-induced aggression (lower panel). SNPs are ordered according to their genomic position. Lead SNPs are marked with an asterisk*. **c,** First upper panel - Kaplan–Meier survival curves for *MAGED1* rs7886851-A, which is associated with the transition from first cocaine use to CUD. Log-rank test, **p value = 0.002. Second upper panel - Kaplan–Meier survival curves for *MAGED1* rs6614508-A, which is associated with the transition from first cocaine use to CUD. Log-rank test, **p value = 0.002. First lower panel – boxplot showing *USP7* rs12446999-A with aggression symptoms. N =237. Wilcoxon–Mann–Whitney *p value = 0.028. Second lower panel - boxplot showing *USP7* rs7342770 with aggression symptoms. N =237. Wilcoxon-Mann– Whitney **p value = 0.0014. Adenosine (A) is the alternate allele. AAO, age at onset; CUD, cocaine use disorder; SAPS, Scale for Assessment of Positive Symptoms – Cocaine-Induced Psychosis.

## Discussion

Here we demonstrated that the major effect of Maged1 on drug-adaptive behavior is mediated outside of the mesocorticolimbic pathway through vGluT2 neurons in the PVT. Furthermore, we showed that the role played by Maged1 is bidirectional in the PVT of adult mice, with significant abolition and restoration of cocaine sensitization. Taking advantage of this phenotype, we found that *Maged1* inactivation significantly altered cocaine-evoked gene repression in the thalamus and further identified its involvement in PRC1 complex activity. The identification of USP7, a well-known PRC1 regulator, and H2Aub as Maged1-interacting partners, together with the observation that chronic cocaine exposure leads to an overall increase in H2Aub+ promoters, indicated that Maged1 mediates cocaine-dependent gene downregulation by controlling histone H2A monoubiquitination. Furthermore, inactivation of *Maged1* in vGluT2 neurons or inhibition of USP7 alters H2A monoubiquitination in the thalamus and cocaine-adaptive behavior. Considering that Maged1 enabled interactions between H2Aub and USP7, the Maged1-USP7 interaction likely constitutes a molecular switch linking PRC1-mediated gene repression to upstream signaling events^31^.

This work also provides a genome-wide assessment of H2Aub deposition and transcriptional alterations in the thalamus in response to repeated cocaine administration. A striking finding of the present study is that, in this non-canonical region, the predominant effect of cocaine is gene repression, conversely to what has been observed in the NAc or the VTA^8^.

Moreover, we observed a significant overlap between cocaine-induced repressed genes and cocaine-induced increase in H2Aub on promoter, further demonstrating that this epigenetic mechanism has an overall significant impact on gene transcription level, while controlling the predominant transcriptomic effect.

Even though evidences suggest that PRC1 complexes are targeted by PTMs, it is still unclear how H2Aub is regulated^42^. Here, we observed that among genes showing an increased H2Aub mark on their promoter and a significant repression induced by cocaine, at least 2 genes, HOPX and SCML, are linked to the PRC1 complexes and may thus represent a regulatory feedback loop.

Our proposed model (Extended Data Fig. 12) postulates that when cocaine is administered, Maged1-USP7 is recruited to H2A, activating or stabilizing PRC1 and leading to H2A monoubiquitination. Thus, in *Maged1*-cKO mice or after treatment with the USP7 inhibitor P5091, PRC1 activity is blocked, leaving it in an unresponsive state such that upstream events (such as cocaine exposure) do not induce its activity (Extended Data Fig. 12). This novel epigenetic mechanism highlighted in mice might be a first step toward developing CUD in human. Indeed, SNP mutations in USP7 and MAGED1 polydrug users led to aggressive behavior during cocaine use and a modified transition time to cocaine addiction, respectively. Although no medication has been developed to be an efficacious treatment for CUD, a chronic disease with harmful consequences worldwide^43,44^, the finding that a pharmacological inhibitor of USP7 is behaviorally effective for a cocaine-relevant behavior in mice might pave the way for the development of new treatments for CUD. The lack of characterization of small-molecule inhibitors and their targeted deubiquitinases is a limiting factor in the development of efficient and selective deubiquitinase inhibitors that could be used in human patients^45^. However, this is now likely to change with new reports of selective USP7 inhibitors that possess a structurally defined mechanism of inhibition^46,47^.

In conclusion, these findings establish a critical role for histone H2A monoubiquitination and its subsequent transcriptional changes, in regulating cocaine-adaptive behaviors, which may constitute an essential epigenetic risk factor for drug addiction in human. Taken together, our ChIP, RNAseq and human genomic analyses clearly position H2Aub in the drug-related epigenetic landscape and link, for the first time, this histone mark to an acquired human disease. The regulation of this newly identified cocaine-induced epigenetic modification might be considered a new potential therapeutic target for prevention and harm reduction in active cocaine users.

## Acknowledgments

We thank Dr. Christian Luscher, Dr. Antoine Adamantidis, Dr. Ami Citri, Dr. Marina Picciotto, Dr. Gilbert Vassart and Dr. Michel George for their critical reading and their helpful insights on our manuscript. We thank Delphine Houtteman, Souad Laghmari, and Laetitia Cuvelier for technical assistance and mice colonies in Brussels. We thank Sarah Gharbi for technical assistance in Grenoble. We thank the EMBL Proteomics Core Facility. JC is a research fellow of FRS-FNRS and Fonds Erasme. LB is an a Ramon y Cajal fellow (CBMSO). AKE is a research director of the FRS-FNRS and a WELBIO investigator.

## Funding

FRS-FNRS grants 23587797, 33659288, 33659296 (AKE)

WELBIO grant 30256053 (AKE)

Fondation Simone et Pierre Clerdent 2018 Prize (AKE)

Fondation ULB (AKE)

Dotation Jeune Chercheur INSERM (LB)

AFM stratégique 2 MyoNeurAlp (LB)

Agence Nationale pour la Recherche grant ANR-21-CE11-0013 (SB)

EMBL Interdisciplinary Postdocs (EIPOD4) initiative co-funded by Marie Skłodowska-Curie (grant 847543) (TC)

Mission Inter-Ministérielle de Lutte Contre la Drogue et la Toxicomanie grant ASE07082KSA (FV)

Direction de la Recherche Clinique et du Développement de l’Assistance Publique − Hôpitaux de Paris grant OST07013 (FV)

French Ministry of Health, Programme Hospitalier de Recherche Clinique National AOM10165 (FV)

## Author contributions

Conceptualization: JC, AKE

Investigation: JC, LB, PH, RI, TC, CD, RI, FV, JB, EM, RC

Visualization: JC, LB, PH, RI, TC, FV, JB, AKE

Data Analysis: JC, LB, RI, MD, TC, SB, FV, EM, RM, AKE

Funding acquisition: SB, FV, AKE

Project administration: JC, AKE Supervision: AKE

Writing – original draft: JC, AKE

Writing – review & editing: JC, LB, RI, MD, TC, SB, FV, JB, AKE

## Data and materials availability

The data supporting the findings are available within the article and its supplementary materials.

## Supplementary Materials

Supplementary Text

Extended Data Figures 1 to 16

Extended Data Tables 1 to 8

References (*48*–*91*)

## Methods

### Animals

The Maged1-inactivated (Maged1 KO) and conditioned (Maged1 lox^p^) mice used in the present study have been previously generated and described^48^.

Maged1 KO do not show gross abnormalities and have normal general cognitive and sensori-motor abilities^9,48–50^. All our experiments were performed in 2-to 4-month-old mice. Our experiments were conducted on hemizygous Maged1 KO males and their wild-type littermates (Maged1 WT) generated by crossing heterozygous Maged1 KO/WT females with C57Bl6J males as Maged1 KO males are deficient in sexual behavior and do not reproduce properly^50^. Conditional knockout mice were generated by crossing heterozygous Maged1 lox^p^/WT or homozygous Maged1 lox^p^/lox^p^ females with different population selective Cre male mice. The Cre-mediated deletion of Maged1 was obtained using mice expressing Cre-recombinase under the control of the vGluT2 promoter, vGluT1 promoter and EMX1 promoter. As with the Maged1 KO mice, we used hemizygous Maged1 cKO males and their floxed littermates to maintain the same conditions allowing comparison of the KO and cKO types. vGluT1cre and vGluT2cre mice were obtained from the Jackson laboratory: Vglut1-IRES2-Cre-D, Stock No: 023527, Vglut2-ires-cre Stock No: 016963. EMX1cre mice^51^. All mice were backcrossed for at least eight generations in the C57Bl6J genetic background. Mice were housed by groups of 2–4 mice, in a 12 h light/dark cycle, with food and water available ad libitum. Experimental procedures were performed in accordance with the Animal Care Committee guidelines and approved by the approved by the ethical committee of the Faculty of Medicine of the Free University of Brussels.

### Genotyping

We systematically check the genotype of each animal by PCR performed on DNA extracted from ear biopsies (1 2 mm). List of the primers used for the wild-type Maged1 and Maged1 KO and lox^p^ alleles are found in de Backer et al.^9^

For vGluT1cre allele, the primers vGluT1mutant CCC-TAG-GAA-TGC-TCG-TCA-AG, common ATG-AGC-GAG-GAG-AAG-TGT-GG, give an amplicon of 344 bp

For vGlut2cre allele, the primers vGluT2mutant (ATC-GAC-CGG-TAA-TGC-AGG-CAA) and common (CGG-TAC-CAC-CAA-ATC-TTA-CGG) give an amplicon of 600 bp

Emx1 : mutant (TCG-ATA-AGC-CAG-GGG-TTC) common (CAA-CGG-GGA-GGA-CAT-TGA) give an amplicon of 195 bp.

### AAV stereotaxic injections

To inactivate Maged1 in a region-specific manner we used Maged1 lox^p^ mice in which we stereotaxically injected an AAV containing the Cre-recombinase in the ventral subiculum and in the PVT (from bregma, in mm): ventral subiculum, A-P -3,64, M-L +-3,15, D-V -4,72 (400 nl), PVT, A-P -1.46, M-L +-1.02, D-V -3.05, (500 nl) angle : 10°.

Behavior experiments after stereotaxic injection aiming at Maged1 inactivation were controlled by AAV-eGFP injection in littermate mice. In specified cases, control mice were Maged1 WT mice injected with AAV-eGFP-cre.

Mice were anaesthetized with avertin (2,2,2-tribromoethanol 1.25%; 2-methyl-2-butanol 0.78%, 20 ll/g, ip, Sigma-Aldrich) before bilateral injection of AAV on a stereotaxic frame (David Kopf Instruments). Injections were performed using a blunt needle connected to a 10μL syringe (Hamilton) and a microstereotaxic injection system (KD Scientific). The injection rate was fixed at 0.1 μL/min. The needle was left in place for 15 min after the injection before being slowly removed. To avoid bias, littermates were equally distributed between the two groups. After surgery, mice were allowed to recover for at least 3 weeks.

We used serotype 5 known for its neuronal tropism, and lowest abilities to migrate anterogradely^52^. Vectors used in this study were purchased from Penn Vector Core (University of Pennsylvania):

- AAV5.CMV.HI.eGFP-Cre.WPRE.SV40 (qPCR-based genome copy titer : 1.864e12)
- AAV5.CMV.PI.eGFP.WPRE.bGH (qPCR-based genome copy titer : 1.704e12)

To re-express Maged1 in a region-specific manner we used full Maged1 KO mice in which we stereotaxically injected an AAV containing Maged1 and an eGFP (linked with a P2A sequence, a 2A self-cleaving peptides inducing a ribosomal skipping during translation), either in the PFC, in the amygdala (PFC, A-P 1.9, M-L +-0.3, D-V -2.3 (600 nl) ; amygdala, A-P -1.0, M-L +-3.0, D-V -4.9 (600 nl)); or in the PVT. The vectors were generated by transient transfection (with X-tremeGENE) of HEK293 cells using three plasmids: our custom-made plasmid, cis ITR-containing CMV-eGFP-P2A-Maged1 (or just CMV-eGFP), the transplasmid encoding AAV replicase and capsid genes (pAAV.DJ/8) and the adenoviral helper plasmid followed by purification (with freezing and thawing and centrifugation) and qPCR-based genome copy titer quantification.

### Cocaine-induced locomotor sensitization

Experiments were performed during the light period of the cycle. To evaluate spontaneous locomotor activity, mice were allowed to freely explore a wide open-field (OF; 40×40×40 cm), made of grey (wall) and white (flour) hard plastic, for 30 to 60 min (as specified). Locomotor activity was recorded in a low luminosity environment (18-22 lx) with a video camera placed on the ceiling above the arena and connected to an automated video tracking system (Ethovision XT 13-14, Noldus Information Technology). Classical cocaine-induced locomotor sensitization: the three-first days of experiment, mice were injected with saline and their locomotor activity was recorded, the next 5 days, mice were injected with cocaine instead of saline (20 mg/kg, intraperitoneal (i.p.)). When specified, we performed a two-injection protocol (15 mg/kg, i.p.) with 3 days of habituation and 7 days between the two cocaine injections, based on Valjent et al. work^15^.

### Cocaine-induced CPP and operant self-administration

Conditioned place preference was conducted in a three chambers apparatus, consisting of a small middle neutral (6 × 20 cm) area connected to two large compartments (18 × 20 cm) that differed in wall and floor conditions (Panlab). Three days before the experiment, mice were used to be manipulated and received a single saline injection in their home cage. At day 0 (preconditioning test, 18 min), mice were placed in the central neutral area and allowed to explore freely both chambers. Mice were randomly assigned to the various experimental groups (unbiased protocol). Conditioning (days 1–6) was carried out as follows: mice were confined to one compartment for 30 min immediately after injection of cocaine (10 mg/kg, ip, or saline for control groups) on days 1, 3 and 5 and to the other compartment after saline injection on days 2, 4 and 6. For the post-conditioning test (day 7), mice were placed in the central area and allowed to explore freely both chambers for 18 min.

For operant self-administration, mice were first anesthetized by a mixture of ketamine hydrochloride (Imalgène; Merial Laboratorios S.A., Barcelona, Spain; 75 mg/kg) and medetomidine hydrochloride (Domtor; Esteve, Barcelona, Spain; 1 mg/kg) dissolved in sterile 0.9% physiological saline. This anesthetic mixture was administered intraperitoneally in an injection volume of 10 ml/kg of body weight. After surgery, a subcutaneous injection of atipamezole hydrochloride (Revertor; Virbac, Barcelona, Spain; 2.5 mg/kg of body weight) was administered to reverse the anesthesia. Additionally, mice received a subcutaneous injection of meloxicam (Metacam; Boehringer Ingelheim, Rhein, Germany; 2 mg/kg of body weight) and an intraperitoneal injection of gentamicin (Genta-Gobens; Laboratorios Normon, S.A., Madrid, Spain; 1 mg/kg of body weight) all previously dissolved in sterile 0.9% physiological saline. For operant conditioning maintained by cocaine, cocaine hydrochloride (Sigma-Aldrich, Saint Louis, MO, USA) was dissolved in a saline solution (0.9% NaCl w/v). The catheter’s patency was evaluated by thiopental sodium (5 mg/ml) (Braun Medical S.A, Barcelona, Spain) dissolved in distilled water and delivered by infusion of 0.1 ml through the intravenous catheter. The experimental sequence was the following; mice were trained under a fixed ratio 1 schedule of reinforcement (FR1; one nose-poke produce the delivery of one dose of cocaine) during 5 consecutive sessions (sessions 1-5) followed by 5 sessions (sessions 6-10) under an FR3 schedule (three nose-pokes produce the delivery of one dose of cocaine).

### Immunohistochemistry

Mice were transcardially perfused with a 0.01 M PBS solution followed by paraformaldehyde 4% in 0.1 M phosphate buffer (pH 7.4). Brains were removed, fixed overnight in paraformaldehyde 4% and successively cryoprotected in 20 and 30% sucrose in 0.1 M phosphate buffer. 20-um-thick coronal sections were cut on a cryostat (-22°). Sections were incubated in 10% normal horse serum in 0.01 M PBS-0.1% Triton X-100 for 60 min at room temperature to block the nonspecific antibody binding. Sections were then incubated overnight at 4°C with the following required primary antibodies: chicken GFP (1:2000, Abcam, ab13970), rabbit mono-ubiquitylated histone H2A (1:1500, Cell Signaling, 8240S), goat USP7 (1/2000, Bethyl Laboratories, A303-943A), mouse mono-ubiquitylated histone H2A (1:75, Merck, 32160702), Maged1(1/300, Cosmo-Bio, BAM-74-112-EX), in 1% normal horse serum, 0.01 m PBS and 0.1% Triton X-100.

After three washes with PBS/0.1% Triton X-100, slices were incubated in PBS for 1 hour at room temperature and incubated 2 hours at room temperature with the appropriated Alexa488 (1/400, Invitrogen), Alexa594 (1/400, Invitrogen) Alexa647 (1/400, Invitrogen) secondary antibodies. Hoechst 33342 (1/5000, H3570, Invitrogen) was used (5 min and washed twice in PBS) for nuclear staining for 5 min.

The sections were next mounted on a Superfrost slide (Thermo Scientific) and dried using a brush before adding Glycergel mounting medium (Dako). Imaging was performed using a Zeiss LSM780 confocal microscope, an AxioImager Z1 (Zeiss) and a wide-field Fluorescent Microscope: Zeiss AxioZoom .v16 all controlled by the Zen Black software (Zeiss).

### In situ hybridization autoradiography

Mice were deeply anesthetized using halothane and killed by dislocation and decapitation, and the whole brain was rapidly removed and frozen on dry ice. 18-um-thick coronal sections were prepared with a cryostat (CM 3050, Leica), mounted onto RNase-free glass-slides (SuperfrostPlus, Thermo Scientific) and stored at -20°C before experiment. In situ hybridization experiments were performed as previously described^9,53^.

### Fluorescent In situ hybridization with RNAscope

RNAscope ISH was used to detect cell type-specific expression of vGluT1 mRNA, vGluT2 mRNA, and Maged1 mRNA. Mice were deeply anesthetized, and the whole brain was removed and rapidly frozen on dry ice. Fresh-frozen tissue sections (16 μm thick) were mounted on positively charged microscopic glass slides (Fisher Scientific) and stored at −80 °C until RNAscope ISH assays were performed. Multiple target gene-specific RNAscope probes were used to observe the cellular distributions of vGluT1, vGluT2 and Maged1 in the whole brain by using a vGluT2 RNAscope probe (Mm-Slc17a6-C2-Vglut-2 probe, cat. no. 319171-C2, targeting 1986–2998 bp of the Mus musculus VgluT2 mRNA sequence, NM_080853.3), a custom-made Maged1 RNAscope probe (Mm-Maged1-O1, cat. no. 563041, targeting 901-1792 bp (targeting exon 4-12) of the Mus musculus Maged1 mRNA sequence, NM_019791.3), VgluT1 RNAscope probe (Mm-Slc17a7-C3-Vglut1 probe, cat. no. 416631-C3, targeting 464 - 1415 bp of the Mus musculus VgluT1 mRNA sequence, NM_182993.2). TSA fluorophores, cyanine3 (NEL744001KT), cyanine5 (NEL745001KT) and fluorosceine (NEL741001KT), were obtained separately at PerkinElmer. All these probes were designed and provided by Advanced Cell Diagnostics. The RNAscope mRNA assays were performed following the manufacturer’s protocols. Stained slides were coverslipped with fluorescent mounting medium (ProLong Gold anti-fade reagent P36930; Life Technologies) and imaged with an AxioImager Z1 (Zeiss) microscope at 20-40x magnification with the use of Zen Black software (Zeiss). Cells expressing vGluT1 mRNA, vGluT2 mRNA, Maged1 mRNA in the cortex (PFC, motor (M1) and somato-sensorial cortex) and the PVT were counted in eight stereologically selected sections per brain under 40x magnification.

### Flow cytometry and cell sorting

Animals were deeply anesthetized using halothane and killed by dislocation and decapitation. Brains were rapidly removed and immersed in ice-cold PBS where whole-thalamus were dissected^54,55^. We then performed a manual dissociation in iced-cold dissociation medium (98 mM Na2SO4, 30 mM K2SO4, 5.8 mM MgCl2, 0.25nmM CaCl2, 1 mM HEPES pH 7.4, 20 mM glucose, 125 uM NaOH)^56^ and filtered the cells through a 40 um nylon mesh right before sorting. Cell sorting was performed using a FACS Aria III cell sorter supported by the FACSDiva software (BD Biosciences). Cells were selected by forward scatter, side scatter, Hoechst dye exclusion and red fluorescence. Cell sorting was carried out at the purity mode of sort precision using a low sheath pressure with a 70 μm nozzle and sorted cells were directly collected into ice-cold phenol and guanidine thiocyanate (QIAzol, QIAGEN) for subsequent RNA extraction. For the assessment of sort quality, cells were sorted in ice-cold PBS and re-analyzed. Total RNA extraction and DNase treatment were carried out using miRNeasy Micro Kit and RNase-free DNase set according to the manufacturer’s instructions (QIAGEN).

### Transcriptome analyses

Total mRNA from six sorted independent samples (3 vGluT2 Cre, loxp-stop-loxp-tdTomato, Maged1 lox^p^ and 3 vGluT2 Cre, loxp-stop-loxp-tdTomato, Maged1 WT) and from 15 unsorted independent samples (vGluT2 Cre::Maged1 loxp or WT, 8 with chronic cocaine, 7 with saline inejctions) was extracted using the QIAGEN miRNeasy micro kit according to the manufacturer’s recommendations. A high vGluT2 neurons proportion invited us to skip the FACS procedure for further transcriptomic experiment (Fig. 2) to avoid any transcriptomic modification relative to the sorting process, and we immediately mechanically dissociated the cells in phenol and guanidine thiocyanate (QIAzol, QIAGEN) which facilitate lysis of fatty tissues and inhibit RNases. Following RNA quality control assessed on a Bioanalyzer 2100 (Agilent technologies), the indexed cDNA libraries were prepared using the NEBNext rRNA Depletion Kit (New England BioLabs), followed by NEBNext Ultra II Directional RNA Library Prep, following manufacturer’s recommendations. The multiplexed libraries (18 pM) were loaded on flow cells and sequenced with a 20 million reads depth, 2×100 bases paired-end (Novaseq 6000, Illumina).

Approximately 20 millions of paired-end reads per sample were mapped against the mouse reference genome (GRCm38.p4/mm10) using STAR 2.5.3a software^57^ to generate read alignments for each sample. Annotations Mus_musculus.GRCm38.87.gtf were obtained from ftp.Ensembl.org. Read counts and transcript per million reads (TPMs) were generated using tximport R package version 1.10.0 and lengthScaledTPM method (Soneson et al., 2016) with inputs of transcript quantifications from tool salmon (Patro et al., 2017). Low expressed transcripts and genes were filtered based on analysing the data mean-variance trend. The expected decreasing trend between data mean and variance was observed when expressed transcripts were determined as which had ≥ 3 of the 40 samples with count per million reads (CPM) ≥ 1, which provided an optimal filter of low expression. A gene was expressed if any of its transcripts with the above criteria was expressed. The TMM method was used to normalise the gene and transcript read counts to *lag*_2_-CPM (Bullard et al., 2010). The principal component analysis (PCA) plot showed the RNA-seq data did not have distinct batch effects. Limma R package was used for 3D expression comparison (Ritchie et al., 2015; Law et al., 2014). To compare the expression changes between conditions of experimental design, the contrast groups were set as Ctrl.Saline-Ctrl.Cocaine, KO.Saline-KO.Cocaine, Ctrl.Saline-KO.Saline, Ctrl.Cocaine-KO.Cocaine, (Ctrl.Cocaine-Ctrl.Saline)-(KO.Cocaine-KO.Saline). For DE genes/transcripts, the *lag*_2_fold change (*L*_2_*FC*) of gene/transcript abundance were calculated based on contrast groups and significance of expression changes were determined using t-test. P-values of multiple testing were adjusted with BH to correct false discovery rate (FDR) (Benjamini and Yekutieli, 2001). A gene was significantly DE in a contrast group if it had adjusted p-value < 0.05.

### Protein extraction

To understand by which means Maged1 modifies genes that are targeted by PRC, we tried to identify its protein partners in saline versus chronic cocaine conditions. On the fifth day of cocaine (or saline) injection, 5 minutes after injection, mice were deeply anesthetized using halothane and killed by dislocation and decapitation. Brains were rapidly removed and immersed in ice-cold PBS where PFC, NAc, and whole-thalamus were dissected. The tissue was mechanically dissociated using an electric rotating pestle (Ultra-Turrax, Merck) and cells were lysed in lysis buffer (before IP-MS and parallel WB: 50mM tris HCL pH 8, 150mM NaCl, 1mM EDTA, 0,5% NP-40, ; for subsequent IP-WB focusing on histones : 10mM tris HCl pH 8, 300mM sucrose, 300mM NaCl, 0,5% NP40) for 25 minutes on ice with protease inhibitor (Complete, Roche, diluted as recommended by the manufacturer) and phosphatase inhibitor (PhosSTOP, Roche, diluted as recommended by the manufacturer). For IP-MS, after a centrifugation at 4°C, 20 minutes, at 15000 g, only the supernatant was used. For histones related subsequent IP-WB, centrifugation at 4°C at 1200 g for 5 minutes separates soluble cytosolic and nuclear proteins (in the supernatant) from chromatin-linked protein (in the pellet). The pellet was treated with a nuclease (Benzonase, Merck, cat. no. E1014) to release mononucleosomes^58^ and related protein-interactions. Proteins were quantified using a Bradford assay (ThermoFisher, cat. no. 23246).

### Immunoprecipitation with mouse lysates

30 ul of Dynabeads Protein A (Invitrogen, cat. no. 10001D) were washed 4 times in protein extraction lysis buffer (without protease or phosphatase inhibitors). The beads were then incubated with the IP antibody (2 ul of anti-Maged1, 7 ul of H2Aub and 7 ul of USP7 were used per sample) for 2 hours at room temperature. The beads were subsequently washed 4 times before proteins were added (500 ug of proteins was added for IP-MS, 1500 ug was used for IP-WB). Samples were incubated overnight with rotation at 4C.

For MS purpose: Beads were washed by the lysis buffer one time, and 3 times in trypsin digestion buffer (20 mM Tris.HCl pH 8.0, 2 mM CaCl2) and resuspended in 150 microliters trypsin digestion buffer.

For WB purposes: Beads were washed by the lysis buffer four times before being resuspended in 40 ul of 2× SDS buffer and thawed 15 minutes at 75°C. The beads were then removed using the magnetic rack.

### Mass spectrometry from mouse lysates

Liquid chromatography–tandem mass spectrometry was done in Francis Impens Lab (VIB Center for Medical Biotechnology, UGent). After pull down, the washed beads were re-suspended in 150 µl trypsin digestion buffer and incubated for 4 hours with 1 µg trypsin (Promega) at 37 °C. Beads were removed, another 1 µg of trypsin was added and proteins were further digested overnight at 37 °C. Peptides were purified on SampliQ SPE C18 cartridges (Agilent), dried and re-dissolved in 20 µl loading solvent A (0.1% trifluoroacetic acid in water/acetonitrile (ACN) (98:2, v/v)) of which 2 µl was injected for LC-MS/MS analysis on an Ultimate 3000 RSLC nano LC (Thermo Fisher Scientific, Bremen, Germany) in-line connected to a Q Exactive mass spectrometer (Thermo Fisher Scientific). Peptides were first loaded on a trapping column made in-house, 100 μm internal diameter (I.D.) × 20 mm, 5 μm beads C18 Reprosil-HD, Dr. Maisch, Ammerbuch-Entringen, Germany) and after flushing from the trapping column the peptides were separated on a 50 cm µPAC™ column with C18-endcapped functionality (Pharmafluidics, Belgium) kept at a constant temperature of 35°C. Peptides were eluted by a stepped gradient from 98% solvent A’ (0.1% formic acid in water) to 30% solvent B′ (0.1% formic acid in water/acetonitrile, 20/80 (v/v)) in 75 min up to 50% solvent B′ in 25 min at a flow rate of 300 nL/min, followed by a 5 min wash reaching 99% solvent B’. The mass spectrometer was operated in data-dependent, positive ionization mode, automatically switching between MS and MS/MS acquisition for the 5 most abundant peaks in a given MS spectrum. The source voltage was 3.0 kV, and the capillary temperature was 275°C. One MS1 scan (m/z 400−2,000, AGC target 3 ×E6 ions, maximum ion injection time 80 ms), acquired at a resolution of 70,000 (at 200 m/z), was followed by up to 5 tandem MS scans (resolution 17,500 at 200 m/z) of the most intense ions fulfilling predefined selection criteria (AGC target 50.000 ions, maximum ion injection time 80 ms, isolation window 2 Da, fixed first mass 140 m/z, spectrum data type: centroid, intensity threshold 1.3xE4, exclusion of unassigned, 1, 5-8, >8 positively charged precursors, peptide match preferred, exclude isotopes on, dynamic exclusion time 12 s). The HCD collision energy was set to 25% Normalized Collision Energy and the polydimethylcyclosiloxane background ion at 445.120025 Da was used for internal calibration (lock mass). QCloud was used to control instrument longitudinal performance during the project^59^.

Analysis of the mass spectrometry data was performed in MaxQuant (version 1.6.11.0) with mainly default search settings including a FDR set at 1% on PSM, peptide and protein level. Spectra were searched against the Mouse proteins in the Uniprot/Swiss-Prot reference database (release version of June 2019 containing 22,282 mouse protein sequences, UP000000589, downloaded from http://www.uniprot.org), supplemented with the protein A recombinant staphylococcus aureus protein sequence. The mass tolerance for precursor and fragment ions was set to 4.5 and 20 ppm, respectively, during the main search. Enzyme specificity was set as C-terminal to arginine and lysine, also allowing cleavage at proline bonds with a maximum of two missed cleavages. Variable modifications were set to oxidation of methionine residues, acetylation of protein N-termini. Matching between runs was enabled with a matching time window of 0.7 minutes and an alignment time window of 20 minutes. Only proteins with at least one unique or razor peptide were retained. Proteins were quantified by the MaxLFQ algorithm integrated in the MaxQuant software. A minimum ratio count of two unique and razor peptides was required for protein quantification. Further data analysis was performed with the Perseus software (version 1.6.2.1) after loading the proteingroups file from MaxQuant.

### Western blot

Samples (input (eluted in 2× SDS buffer and thawed 15 minutes at 75°C) and IP proteins) were run in NuPAGE 4%–12% Bis-Tris Mini Protein Gel (10 wells) (ThermoFisher Scientific, NP0321BOX) at the voltage of 100V for 2.5 hours in a NuPAGE MOPS SDS Running buffer (ThermoFisher, cat. no. NP0001) and then transferred to Nitrocellulose Blotting Membrane (Bio-Rad, cat. no. 1704270, 0.2um) and placed in a Trans-Blot Turbo Transfer System (Bio-Rad) (1.3 A constant and up to 25 V) for 10 minutes. PageRuler Plus Prestained Protein Ladder, 10 to 250 kDa was systematically used (ThermoFisher, cat. no: 26619). The membrane was blocked in the intercept TBS blocking buffer (Li-Cor, cat. no. 927-60001) buffer for 1 hour at room temperature and subsequently incubated in the blocking buffer containing rabbit anti-Maged1, (1/3000, Cosmo-Bio, BAM-74-112-EX), goat anti-USP7 (1/700, Bethyl Laboratories, A303-943A), rabbit anti-H2A (1/1000, Merck, 07-146), and rabbit anti-mono-ubiquitylated H2A (1:1000, Cell Signaling, 8240S) antibodies overnight at 4C.

For qualitative analysis, this was followed by the incubation in the blocking solution containing secondary antibody anti-Rabbit or/and goat IgG antibody conjugated with HRP (GENA934, AP106P, Merck) at room temperature for 1 hour for enhanced chemiluminescence. Pierce ECL Western Blotting Substrate (Cat. #32106, ThermoFisher Scientific) was used for signal detection. For quantitative purposes, this was followed by the incubation in the blocking solution containing secondary antibody goat anti-Rabbit DyLight 680 (ThermoFisher, cat. no. 35568) or/and donkey anti-Goat DyLight 680 (ThermoFisher, cat. no. SA5-10090). For quantification, ImageStudioLite version 5.2.5 (LI-COR, 2014) was used. We defined regions of interest (ROIs) for each band of interest and performed background subtraction to remove any non-specific signals. We then measured the intensity of each band within the ROIs. We normalized the band intensities to the appropriate reference protein. We then imported the quantification data into a GraphPad Prism and performed two-way ANOVA to compare the different conditions.

The raw representative WB gel and immunoblot following IP are shown in Extended Data 15.

### Human cell lines

Human embryonic kidney 293T (HEK293T) and human neuroblastoma cells (SH-SY5Y) were cultured in Dulbecco modified Eagle’s medium containing 10% fetal bovine serum (FBS Qualified, Gibco) at 37°C in an atmosphere of 5% CO2.

The Maged1 KO in HEK cells was generated by using the RNA-guided CRISPR-Cas nuclease system^60^. For the design the Synthego and E-CRISP bioinformatic tools were used to minimize off-target effects. The input target genomic DNA sequence used was the human genome GRCh38.p13 from ENSEMBL where Maged1 gene locates on ChrX p11.22. As a results of the analysis six 20-nucleotide guide sequence were selected between exon 2 and exon 5 (gRNA1=GGCAGGGCGTTATACTACAG, gRNA2=CAAGGCGCTGTCTTCTACG, gRNA3=GCCTCAGCCTGCAAACACAG, gRNA4=AATGTTGAAGAGAACAGCAG, gRNA5=GTGGAATTGGCCAATCAGCG, gRNA6=GATGCTTAAGGACTACACAA). The 6 gRNAs were synthetized inside a U6RNA cassette between the U6 promoter and the gRNA scaffold by Twist Bioscience. Starting from the 6 cassettes nine different combinations of tandem gRNA were coupled in order to cut at the extremities of exon 2 and exon 5 (combo 1= gRNA1+4, combo 2= gRNA1+5, combo 3= gRNA1+6, combo 4= gRNA2+4, combo 5= gRNA2+5, combo 6= gRNA2+6, combo7= gRNA3+4, combo 8= gRNA3+5, combo 9= gRNA3+6) and directly transfected in HEK 293T cells with an empty Cas9 plasmid. HEK 293FT cells were maintained according to the manufacturer’s recommendations. Cells were cultured in DMEM medium supplemented with 10% (vol/vol) FBS at 37 °C and 5% CO2. The cells were plated onto 6-well plates without antibiotics 16-24h before transfection and seeded at a density of 0.5 × 10^6^ per well in a total volume of 2 ml. Cells were transfected with polyethylenimine (PEI) and Cas9 plasmid with the 2 gRNA mixed at equimolar ratios using 1μg of total DNA (0.7 μg of pX459-Cas9-Puro + 96ng for each gRNA), 2 μg/ml of puromycin was applied 24 hours after transfection for 48 hours. The clonal-density dilution procedure was then applied to isolate the clonal cell line and expanded for 2-3 weeks. Finally, the DNA was extracted by using Monarch® Genomic DNA Purification Kit (NEB, T3010S). For the analysis of microdeletions of the targeted exons an Out-Fwd (p1 GCACAGTCAGACCACAGTCACC) and Out-Rev primers (p2, GAGCCAACCATGGAAAGGAGGG) both of which were designed to anneal outside of the deleted region, were used to verify the successful deletion by product size. A parallel set of PCRs with In-Fwd (p1 same as above) and In-Rev primers (p3 CTGCGGTGGCCTGGTTAGTAG) was designed to screen for the presence of the WT allele. By analyzing the PCR result the combo 6 (gRNA 2+6) was chosen as the best combination to delete the region between exon 2 and 5, even if a wild type band is still present. This was expected since HEK293T cells have 3 X chromosomes containing the *MAGED1* gene, so the heterozygous clone selected (clone A1) was re-transfected with Cas9 plasmid + gRNA 2+6, two isolated sub-colonies were obtained and the abolished expression of MAGED1 at protein level was confirmed by WB (Fig. 2k). The sub-clone C1 was selected and used for subsequent experiments.

### Immunoprecipitation from human cells

For MAGED1 co-IP in HEK cells a KO cell line for *MAGED1* was used as control, while in SH-SY5Y cells rabbit IgG-IP was used as isotype control respect to MAGED1 antibody (polyclonal IgG rabbit). Wild-type HEK293T, *MAGED1* KO HEK293T and SH-SY5Y cells were grown in triplicates in 150 mm dishes (20.0 × 106). Cells were lysed with lysis buffer (50 mM Tris–HCl pH 7.5, 150 mM NaCl, 1% Triton X-100) and supplemented with protease inhibitors (Roche, cat.no. 11836170001). Cell lysate was incubated for 30 min at 4°C, sonicated for 5 min at 0°C and then centrifuged at 15000 g for 15 minutes. The total protein concentration was determined by BCA assay and 500 μg of total proteins was incubated for 1 hour (4°C, rotary agitation) with 25 μl of Pierce™ Protein A Agarose beads (ThermoFisher, cat. no. 20333), centrifuged at 2500 g for 3 minutes at 4°C and the supernatant recovered as pre-cleared lysate. For the immune complex formation (total proteins + antibody) 500 μg of the pre-cleared lysate was incubated overnight (4°C, rotary agitation) with 2.5 μg of MAGED1 antibody (ThermoFisher, cat. no. PA5-99091) or 2.5μg of IgG rabbit (Biotechne, cat.no. NB810-56910) as isotype control. To reduce nonspecific binding and background, 25 μl of protein A Agarose beads were incubated for 1hr with 0.1% of BSA (4°C, rotary agitation) and washed two times with 500 μl of IP buffer (50 mM Tris–HCl pH 7.5, 150 mM NaCl). The pre-blocked protein A beads were added to the immune complex at 4°C for 4 hours under rotary agitation, centrifuged at 2500 g for 3 minutes at 4°C to discard the flowthrough and washed three times with 500 μl of IP buffer and one time with PBS only. The immune complex bound to the beads was eluted in 60μl of Laemmli buffer 2X (Biorad cat.no. 1610747) incubated for 10 minutes at 95°C, centrifuged at 2500 g for 3 minutes and sent for MS analysis.

### Mass spectrometry from human cells

Liquid chromatography–tandem mass spectrometry was here performed by the Proteomic Core Facility of EMBL (Heidelberg, Germany).

### Sample preparation

Disulfide bridges were reduced in cysteine using dithiothreitol (56°C, 30 min, 10 mM in 50 mM HEPES, pH 8.5) and such reduced cysteines were then alkylated with 2-chloroacetamide (room temperature, in the dark, 30 min, 20 mM in 50 mM HEPES, pH 8.5). Established protocols ^61,62^ were followed and trypsin (Promega, sequencing grade,) was added (1/50 enzym/protein ratio) for overnight digestion at 37°C. Next day, the peptide recovery was performed by collecting supernatant on magnet and combining with second elution wash of beads with HEPES buffer. Peptides were labelled with TMT6plex^63^ Isobaric Label Reagent (ThermoFisher) according the manufacturer’s instructions. For the TMT6plex samples were combined and for the sample clean up an OASIS® HLB µElution Plate (Waters) was used. Offline high pH reverse phase fractionation was carried out on an Agilent 1200 Infinity high-performance liquid chromatography system, equipped with a Gemini C18 column (3 μm, 110 Å, 100 × 1.0 mm, Phenomenex)^64^.

### LC-MS\MS

An UltiMate 3000 RSLC nano LC system (Dionex) fitted with a trapping cartridge (µ-Precolumn C18 PepMap 100, 5µm, 300 µm i.d. × 5 mm, 100 Å) and an analytical column (nanoEase™ M/Z HSS T3 column 75 µm × 250 mm C18, 1.8 µm, 100 Å, Waters) were used. Trapping was carried out with a constant flow of trapping solution (0.05% trifluoroacetic acid in water) at 30 µL/min onto the trapping column for 6 minutes. Peptides were eluted by the analytical column with solvent A (0.1% formic acid in water, 3% DMSO) with a constant flow of 0.3 µL/min and increasing percentage of solvent B (0.1% formic acid in acetonitrile, 3% DMSO). The outlet of the analytical column was coupled directly to an Orbitrap Fusion™ Lumos™ Tribrid™ Mass Spectrometer (Thermo) using the Nanospray Flex™ ion source in positive ion mode. The peptides were loaded into the Fusion Lumos via a Pico-Tip Emitter 360 µm OD × 20 µm ID; 10 µm tip (New Objective) and a spray voltage of 2.4 kV was applied with capillary temperature of 275°C. Full mass scan (MS1) was acquired with mass range 375-1500 m/z in profile mode in the orbitrap with 60000 resolution. The maximum filling time was of 50 ms. Data dependent acquisition (DDA) was performed with the resolution of the Orbitrap set to 15000, with a fill time of 54 ms and a limitation of 1×105 ions. A normalized collision energy of 36 was applied. MS2 data was acquired in profile mode.

IsobarQuant^65^ and Mascot (v2.2.07) were used to process the acquired data, which was searched against a Uniprot Homo sapiens proteome database (UP000005640) containing common contaminants and reversed sequences. The following modifications were included into the search parameters: Carbamidomethyl (C) and TMT6(K) (fixed modification), Acetyl (Protein N-term), Oxidation (M) and TMT6 (N-term) (variable modifications). For the full scan (MS1) a mass error tolerance of 10 ppm and for MS/MS (MS2) spectra of 0.02 Da was set. Other parameters chosen: Trypsin as protease with an allowance of maximum two missed cleavages: a minimum peptide length of seven amino acids; at least two unique peptides were required for a protein identification. The false discovery rate on peptide and protein level was set to 0.01. The raw output files of IsobarQuant (protein.txt - files) were processed using the R programming language (ISBN 3-900051-07-0). Only proteins that were quantified with at least two unique peptides were considered for the analysis. Raw signal-sums (signal_sum columns) were first cleaned for batch effects using limma^66^ and further normalized using variance stabilization normalization^67^. Proteins were tested for differential expression using the limma package. The replicate information was added as a factor in the design matrix given as an argument to the ‘lmFit’ function of limma. Also, imputed values were given a weight of 0.05 in the ‘lmFit’ function. A protein was annotated as a hit with a false discovery rate (fdr) smaller 5 % and a fold-change of at least 2 and as a candidate with a fdr below 20 % and a fold-change of at least 1.5.

### USP7 inhibition and in vivo cannula infusion

USP7 inhibitor: P5091 (Selleck Chemicals, S7132) was prepared freshly before injection. P5091 was administered (10 mg/kg i.p. or 0.1 mg/kg intrathalamic) every day during the two-injection cocaine sensitization protocol (just before saline or cocaine administration (15 mg/kg) and at day 2-6) (Fig. 3).

For i.p. injection: the powder was first pre-dissolved in 5% v/v DMSO (vortex and sonication were used to aid dissolving and obtain a clear mix) then 95% v/v sesame oil (vortex was used to aid dissolving and obtain a clear mix) was added to the mix. Calculation was done to inject in i.p. a volume of 100 ul maximum and a dose of 10 mg/kg.

For cannula injections: the powder was first pre-dissolved in 54% v/v DMSO (vortex and sonication were used to aid dissolving and obtain a clear mix) then 26% v/v sesame oil (vortex and sonication were used to aid dissolving and obtain a clear mix) followed by 20% of artificial cerebrospinal fluid (NaCl 125 mM, KCl 2.5 mM, NaH2PO4 1.25 mM, NaHCO3 25 mM, MgCl2 1 mM, CaCl2 2 mM, D-glucose 25 mM) was added to the mix. Calculation was done to inject through cannula a volume of 400 nl and a concentration of 7.5 g/l.

We implanted 21-gauge guides that were cut 5.5 mm below a plastic pedestal (Plastics One Inc.) 0.2 mm above the PVT (from bregma, in mm): A-P -1.46, M-L +-1.02, D-V -2.8, angle: 10° and secured on the skull after H2O2 preparation using a dental cement (Metabond kit, Parkell Inc.). For daily injection, an internal cannula of 26-gauge projecting 0.2 mm below the guide was inserted into the guide and connected to a 10μL syringe (Hamilton). The injection rate was fixed at 0.1 μL/min. Contention was limited as the mice could freely move in their home cage during injection. No anaesthetics were administered for daily injections.

### ChIPmentation

ChIPmentation experiments were performed based on the protocol of Schmidl C et al (doi: 10.1038/nmeth.3542) with minor modifications. The thalami of 2-4 WT or *Maged1* cKO mice injected either with cocaine or saline vehicle were microdissected and crosslinked in freshly prepared methanol free 1% formaldehyde solution (Thermofisher #28906) for 10 min at room temperature. The crosslinking reaction was stopped with0,125M Glycine solution and the samples were washed 3x with PBS and sored at -80C. Samples were then resuspended in cell lysis buffer (10 mM Tris pH 8.0, 10mM NaCl, 0,2% NP40, 1× EDTA free Proteinase inhibitor cocktail-PIC; Sigma # 11836170001) and nuclei were released by homogenization with a pestle B –douncer, centrifuged 8 min at 600g (4C) and resuspended in 130ul of sonication buffer (50 mM Tris pH 8.0, 10mM EDTA, 0,25% SDS, 1× PIC). After 15 min at 4C the resuspended nuclei were transferred to Covaris sonication microtubes (Covaris # 520045) and sonicated in a Covaris S220 (to fragment chromatin to an average size of 200-400bp. Sheared chromatin was centrifuged at >13.000g for 10 min to eliminate cell debris, and transferred to a new 1,5ml tubes and diluted 2,5x with ChIP dilution buffer (20mM HEPES pH 7.3, 1mM EDTA, 0,1% NP40, 150mM NaCl, 1× PIC). Anti-H2AK119ub antibody–Dynabeads protein G complexes were added to the diluted chromatin and incubated overnight at 4C with gentle rotation. For that, for each ChIP sample, 25ul of Dynabeads protein G magnetic beads (Thermofisher #) were transferred to 1,5 ml tubes and washed twice with 1 ml of blocking buffer (PBS1x, 5mgr/ml BSA, 0,5% Tween 20, 1XPIC), resuspended in 200ul of blocking buffer and conjugated with 4ugr of anti-H2AK119ub antibody (Cell Signaling, 8240S) for 3h at 4C in a rotating platform. Subsequently beads-antibody complexes were washed twice and resuspended in 15ul of blocking buffer and added to the diluted chromatin. For each wash, the beads were incubated 5 min at 4C on a rotating platform, collected for 2 min in a magnetic stand and resuspended in 1 ml of blocking buffer.

After overnight incubation chromatin –antibody-Dynabead complexes were washed twice with 1ml of RIPA/low salt buffer (10 mM Tris pH 8.0, 140mM NaCl, 0,1%, 1mM EDTA, Sodium Deoxicholate, 0,1% SDS, 0,1% Triton X100, 1XPIC), twice with 1ml of RIPA/high salt buffer (10 mM Tris pH 8.0, 500mM NaCl, 0,1%, 1mM EDTA, 0,1% Sodium Deoxicholate, 0,1% SDS, 0,1% Triton X100, 1XPIC), twice with 1ml of LiCl buffer (10 mM Tris pH 8.0, 250mM LiCl, 1mM EDTA, 0,5% Sodium Deoxicholate, 0,5% NP40, 0,1%, 1XPIC) and twice with 1ml 10mM Tris pH8.0. Washes were performed as described for the beads-antibody conjugation. After the last wash beads were resuspended in 200ul of 10mM Tris pH8.0 and transferred to a new tube, collected with the magnet and resuspended in 24ul of Tagmentation Mix (1:1 Illumina 2× Tagmentation buffer: 10mM Tris pH8.0). Resuspended chromatin-antibody complexes were incubated for 5 min at 37C and 1ul of Illumina TDE1 enzyme was added to each sample. After mixing each ChIP sample was tagmented for exactly 7 min at 37C and the reaction was immediately stopped by the addition of 1 ml of RIPA/low salt buffer. Samples were then washed once more with 1ml of RIPA/low salt buffer and once with 1ml TE buffer (10 mM Tris pH 8.0, 1mM EDTA). Finally, chromatin was eluted by resuspending beads in 100 ul of Elution buffer (10 mM Tris pH 8.0, 5mM EDTA, 0.1% SDS, 0.3M NaCl) supplemented with 50ugr of proteinase K and incubating at 65C overnight in an agitating thermoblock with heated lid. For Total Imput Chromatin (TIC) samples, 10 ul of sonicated chromatin were diluted 10x in elution buffer with 50ugr of proteinase K and decrosslinked at 65C overnight. In all cases, decrosslinked DNA was treated with 20ugr of RNAseA for 10 min at 37C, and purified with the Qiagen minielute PCR purification kit (Qiagen # 28004). Approx 10-20ng of TIC samples were then diluted in Illumina Tagmentation buffer and tagmented with the TDE1 enzyme as for ChIP samples. Finally, DNA was re-purified with the Qiagen minielute PCR purification kit and size-selected with the Pronex paramagnetic beads.

In all cases, 2 ul of each sample were used for library quantification using the KAPA Library Quantification Kit (Roche #KK4923). The rest of the DNA eluate (20ul) was used for library amplification using Nextera single-index primers as described in Schmidl C et al. Using the KAPA HiFi hotstart ready mix (Roche # KK2601). The number of cycles used to amplify each library was determined as the Ct value of the qPCR reaction +1. For library quantification and amplification the KAPA HiFi polymerase was preactivated by heating at 98C for 30 sec. In all cases, a nick repair step was performed before PCR amplification by incubating the PCR reactions at 72c for 6 min. PCR-amplified libraries were purified and size selected for fragments of 200-250bp using the Pronex paramegnetic beads (Promega # NG2001) according to manufacturer instructions. Finally, libraries were quantified using the Qubit High Sensitivity DNA quantification kit (ThermoFisher # Q32851) and analyzed by Fragment analyzer and sequenced pair paired-end sequencing 100bp in a MGISEQ-2000 platform (BGI sequencing services, Honk Kong).

### H2Aub enrichment analysis

ChIPmentation sequencing data were analyzed using the public Galaxy server (https://usegalaxy.eu)^68^ and personalized R scripts. First, adapter sequences were trimmed from both R1 and R2 read files using Cutadpat v3^69^ and mapped against the mouse mm10 genome assembly using Bowtie2 (v 2.3.4.1)^70^. Significantly enriched regions were called using MACS2 (v 2.1.1.20160309.3) with broadpeak calling parameters^71^. The bam files from TIC samples were used as a control for MACS peak calling. Coverage files were normalized by the number of millions of valid paired reads of each sample. The R (v4.1.2) and Rstudio (1.4.1106) softwares were used for downstream analysis.

For the analysis of H2Aub+ elements, only MACS2 peaks that were in common in at least two biological replicates were selected for further analysis using the R GenomicRanges package (v1.44.0)^72^. The distance distribution of peaks from the nearest TSS was calculated using the chipenrich package^73^. The mouse GCR38 gene annotation was obtained using the R annotationHub package (v3.0.2) and used to select gene promoters (defined as the genomic region 5kb upstream and 2kb downstream of the gene TSS) overlapping with H2Aub peaks. Differentially enriched H2Aub+ genes were thus defined as those loci that displayed H2Aub+ promoters between two experimental conditions. We also identified differential H2A enriched genes by comparing H2Aub+ elements called in saline vs cocaine injected mice and attributing these peaks to genes using chipenrich package (polyenrich method). The conclusions of this analysis were overall similar to those reached based on the analysis of H2Aub promoters (data not shown). However, we preferred to base subsequent analyses on H2Aub+ promoter because of the limitations of long-range enhancer assignment to target genes.

The H2Aub antibody used in this study has been used in ChIPseq experiments in a number of previous works (e.g Kallin et al., 2009^74^). The genome-wide distribution of H2Aub enriched regions closely parallels that of other studies^74,75^ with a high enrichment at gene promoters (Extended Data Fig. 16). Besides, this distribution differs from the ubiquitous and promoter enriched distribution described for H2A (e.g, Kallin et al., 2009^76^), supporting that our H2Aub antibody specifically recognize this epigenetic modification rather than non-specifically immunoprecipitating H2A-bound chromatin. As shown in Extended Data Fig. 16, H2A is broadly distributed genome-wide with only mild enrichment over gene bodies and is depleted from the nucleosome free regions of gene transcription start and transcription termination sites. Instead, H2Aub is mostly enriched at gene promoters. Thus, the differences in our H2Aub coverage profiles with those reported for H2A and their resemblance to the H2A distribution of Yang et al., 2014^75^ strongly suggests that the H2Aub enrichments reported in our study are specific.

### Gene Ontology and curated gene set analysis

Gene Ontology analyses of biological processes of genes displaying either altered H2Aub binding in their promoters and/ or that were differentially expressed across the different experimental conditions were performed using the gprofiler2 (v0.2.1), ClusterProfiler (v4.0.5) and enrichplot (v1.12.3) packages^77,78^. For differentially expressed gene sets, gprofiler analysis was performed by ranking genes according to their FC value, while the unranked gprofiler test was used for gene sets displaying differential H2Aub binding in their promoters or for gene lists resulting from the crossing of RNAseq/ ChIPmentation data analysis. Finally, the GSEA software, a joint project of UC San Diego and Broad Institute^79^, was used for curated gene set analysis of differentially expressed genes.

### Statistics

Statistical analyses were performed using GraphPad Prism 9 software (GraphPad Software Inc.) and SPSS statistics 27 (IBM).

Parametric data, provided by the Kolmogorov–Smirnov test, were analysed using a Student’s t-test, one-way and two-way ANOVA tests (when a value was missing (because of a random reason, we performed a mixed-effects model instead of a two-way ANOVA) and Tukey’s or Sidak’s post hoc tests when appropriate, as indicated in figure legend. For nonparametric data, the Wilcoxon Mann-Whitney test for two-sample comparisons or the Kruskal–Wallis test for multiple comparisons were used. Differences were considered significant when P < 0.05. The results are expressed as mean n standard error of mean (s.e.m.). Threshold for significance was set at P < 0.05. Sample sizes (“n”) are given as the number of animals used for experiments.

For RNAseq statistical analysis we used the The 3D RNA-seq. To establish a significant differential list of genes, a FDR < 0.05 was chosen and a log2Fold-change threshold > 0.5 or <0.5 was applied when mentioned.

### Human genotyping

Human genotyping was performed in collaboration with Romain Icick and Florence Vorspan (Université de Paris, Inserm UMR-S1144).

### Sample selection and clinical assessment

Patients > 18 years seeking treatment for any SUD other than nicotine were consecutively recruited between April 2008 and July 2016 through two multicentric protocols. Both protocols and the current study were approved by the relevant Institutional Review Boards [CPP Ile-de-France IV and CEEI from the Institut de la Santé et de la Recherche Médicale (INSERM), IRB00003888 in July 2015, respectively]. The clinical trial registration numbers are: NCT00894452 & NCT01569347All participants provided written informed consent for both the clinical and genetic assessments, and study records were continuously monitored by the hospital research administration. The research was conducted in accordance with the Helsinki Declaration as revised in 1989. Eligible participants were excluded if they had severe cognitive impairment or insufficient mastery of the French language preventing misunderstanding of the study purposes and assessments, if they had no social insurance, and if they were under compulsory treatment or mentorship.

Assessments included history of substance dependence using the E section of the Structured Clinical Interview for DSM-IV (SCID-IV)^80^ and sociodemographic elements (currently married, yes vs. no; currently employed at least 20% time job, yes vs. no, ever been homeless – defined as having spent at least three months in the streets, yes vs. no). For the current study, we focused on cocaine-related variables that were modelled in the preclinical study, namely: lifetime cocaine dependence (presence/absence), age at onset (AAO) of first cocaine use and of cocaine dependence (transition from first to dependent cocaine use (hereafterm termed “TRANSCOC”) - considered to be related to the rewarding properties of cocaine, which were modelled in the preclinical part - and the behavioral components of the Scale for Assessment of Positive Symptoms – Cocaine-Induced Psychosis (SAPS-CIP) considering total score, agression and stereotypies)^81^.

### Biological sampling and genetic analyses

DNA was extracted from whole blood using a Maxwell 16 PROMEGA® extractor (Promega France, Charbonnières-les-Bains, France). Purity assessment followed the procedures described by the Centre National de Recherche en Génomique Humaine. Participants were genotyped using the Infinium PsychArray (Illumina, San Diego, CA, USA) in two batches (2015 and 2017) by Integragen SA® (Evry, France) using the same pipeline. Genotype files were merged for the present study, keeping only bi-allelic variants common to both extractions. Then, PLINK 2.0^82^ was used for quality control (QC), based on a consensus procedure for ancestry, relatedness and genetic discrepancies^83^, followed by imputation on the Michigan Imputation Server v1.0.4 using the reference file hrc.r1.1.2016 through a SHAPEIT + MiniMac3 pipeline^84^. The number of post-QC variants increased from 500,000 to 7,833,448 SNPs. Chromosome X was imputed separately by Impute2 after pre-phasing by Shapeit4, following a consensus procedure^85^, after exclusion of the pseudo-autosomal region. This increased the number of post-QC variants from 12,942 to 34,243. Quality control and ancestry assessment were performed on a whole-genome level using a standard procedure^83^ based on: an overall call rate >99%, missing genotypes per study and per individual <5%, Hardy-Weinberg equilibrium held at p >10-6 and relatedness inferior to the 2nd degree (in case two participants fell within such relatedness, the one with the highest overall call rate was kept for the study). From 592 genotyped patients, 522 were left for analysis, 351 of whom had strict Caucasian ancestry as compared to the distribution of variants in European from the 1000 genomes database.

The candidate gene study was performed using genomic positions (human genome version GChR37) of the genes prioritized by the preclinical study, keeping markers with minor allele frequency (MAF) >5%, namely: *MAGED1* (chromosome X, 72 markers), *USP7* (chromosome 16, 142 markers), and histone 2A genes (chromosome 1, HISTH2AC18 and HISTH2AC20, no marker). Variants showing significant associations with any of the phenotypes of interest were annotated for their biological function and plausible impact according to multiple knowledge-based repositories online, as previously suggested^86^: the Combined Annotation-Dependent Depletion (CADD) database^87^, brain expression and methylation quantitative trait loci (eQTL through BRAINEAC, http://www.braineac.org/, and mQTL through the mQTLdb database, http://www.mqtldb.org), and their ability to bind DNA enzymes and/or modify DNA conformation (regulomeDB database http://www.regulomedb.org). We also considered linkage disequilibrium (LD, using the r2 measure), which indicates how minor alleles tend to be inherited altogether due to chromatin blocks. High LD (considered at r2 >0.9 in our study, based on European reference panels) indicate a similar genomic signal from possibly multiple SNPs, among which both the strongest statistical association and the highest functional impact can be used to identify the one(s) responsible for the biological signal behind the statistical association. Additionally, we identified CNVs following a pipeline^88^ based on the Illumina® final report file generated by GenomeStudio v.2, which involves two programs, PennCNV^89^ and QuantiSNP^90^, in a combined algorithm. Both programs are based on the hidden Markov chain model but show different sensitivity and specificity. CNVs may be very rare, but represent the closest variation to gene deletion in rodent models.

### Data analysis

Continuous variables were described by means (standard deviation, SD) or medians (interquartile range, IQR) depending on their distribution, and qualitative variables are described by absolute counts and frequencies. Nonparametric tests were used to identify the clinical and sociodemographic factors associated with cocaine-related variables.

SNP-based analyses were performed using PLINK linear regression adjusted for sex and the first ten principal components of ancestry, a stringent adjustment recommended to further minimize the risk of population stratification. Possibly censored variables related to time, namely: AAO of cocaine dependence and transition from 1^st^ cocaine use to CUD, which were analyzed using Cox proportional regression models in R. All these analyses included age, biological sex, and the first ten ancestry components as covariates and cofactor. CNVs in the candidate genes -if any - were also tested for associations with cocaine-related variables based on frequency, size or type (duplication *vs*. deletion) in the subsample of Caucasian ancestry. Statistical significance was set at *p* <0.05 after correction for multiple testing using a false discovery rate procedure (Benjamini-Hochberg, BH-FDR). Separately for chromosome X and 16, the *p*-values of all seven phenotypes and SNPs were listed together to yield the corresponding FDR. In the case of multiple statistical associations (which is expected when using imputation), we used linkage disequilibrium (LD, measured by r2) and *in silico* gene expression measures in the brain (http://www.braineac.org/, Extended Data Table 7) to identify lead SNPs as the most plausibly associated with the underlying biological signal. These lead SNPs (identified as having LD r^2^ <0.9, strong statistical association and reliable evidence for functional impact) were further characterized using graphical representations and adjustment for more sociodemographic and clinical confounders, depending on the phenotype they were associated with. Analyses were conducted with R 4.02 through R studio 1.4.1103 under Mac OS X.12.6. The current study follows the STREGA guidelines for the report of genetic studies^91^.

The summary of the R session and the summary statistics (Extended Data Table 8) are available as Supplementary files. Of note, no QQ-plots were produced given the within-cases, candidate-gene design of the study, driven by preclinical experiments.

### DATA availability

The data supporting the findings are available within the article and its Supplementary Materials and are available from the corresponding author upon request. All RNAseq and ChIPmentation data are deposited in GEO under accession number GSE208142. The R scripts used for this study are available in GitHub upon request.

## Extended Data for

### Supplementary Text

#### RNAseq

To assess the proportion of vGluT2 neurons in the dissected thalamus, we performed fluorescence-activated cell sorting (FACS) of vGluT2 neurons specifically expressing tdTomato, and we found that 93.1% of sorted cells showed vGluT2 expression (Extended Data Fig. 6a). A principal component analysis separated the replicates of the microdissected thalamus data on the basis of mouse genotype and cocaine or saline treatment (Extended Data Fig.6b). In control mice, repeated cocaine injection induced upregulated expression of 252 genes and downregulated expression of 452 genes compared to the expression of these genes after saline treatment (Fig. 2b and Extended Data Fig. 6c).

A gene ontology (GO) term enrichment analysis indicated that genes with cocaine-induced downregulated expression were significantly associated with microtubule regulation, axonal projection assembly and ion transport, while genes with cocaine-induced upregulated expression were assigned to supramolecular fiber (actin) organization and apoptosis (Extended Data Fig. 6d). Thus, cocaine-dependent downregulation and upregulation of gene expression seemed to affect different, yet related, biological processes.

Only 145 genes were similarly downregulated by cocaine treatment in mice with the *Maged1*-cKO and control genotypes, and 297 genes were specifically repressed in a cocaine-dependent manner only in the *Maged1*-cKO mice (Fig. 2b). Moreover, the expression of many genes was upregulated by cocaine in Maged1-cKO mice (494 genes), of which the expression of 383 (77.5%) genes was not upregulated in control mice (Fig. 2b). Thus, *Maged1* inactivation mainly led to an altered transcriptional downregulation in the thalamus after cocaine administration (Fig. 2b).

#### Cocaine-adaptive behaviors

To dissociate a global effect on locomotor activity rather than a specific effect on cocaine-induced sensitization of Maged1 in the PVT, we compared the locomotor activity on the first day of saline injection and the last day of cocaine injection (after reaching the ceiling level) in the rescue model, Maged1 KO, AAV-Maged1 (PVT), and in the cKO model Maged1 loxP, AAV-Cre (PVT). While there was no spontaneous locomotor activity difference between the inactivation or the re-expression of Maged1 in the PVT in these 2 models, the sensitization was significantly stronger in the rescue group as illustrated by an interaction effect between mice group (rescue versus deletion in the PVT) and treatment (saline/cocaine) (Extended Data Fig. 13).

The primary reinforcing properties of cocaine were evaluated in vGluT2cre::Maged1loxP mice and corresponding controls using operant self-administration. Mice were first trained in 5 sessions of fixed-ratio 1 (FR1) followed by 5 sessions of FR3 (Extended Data Fig. 3b). During FR1 and FR3 sessions; no statistical differences were obtained in active nose-pokes between genotypes (Extended Data 3a, two-way ANOVA repeated measures, genotype F1,14=1.130, n.s.). The number of nose-pokes on the inactive lever was similarly reduced in both genotypes (Extended Data 3c). No difference was observed in active nose-pokes and number of infusions during the last two or three days of FR1 and FR3, respectively (genotype × time, F9,126=1.529, n.s., two-way ANOVA repeated measures). Similarly, the percentage of mice reaching the criteria of operant conditioning learning was 66.7% for WT mice and 75.0% for vGluT2cre::Maged1loxP mice following FR1 and FR3 training (chi-squared test=0.125, n.s.). Importantly, during the three consecutive days after the acquisition criteria of stability, discrimination and more than ten infusions, the number of active nose-poking responses was significantly higher in the vGluT2cre::Maged1loxP mice than in the control mice (Extended Data 3d).

#### Human data

In the clinical sample, we showed that the associations between MAGED1 and transition from 1st cocaine use to CUD and between USP7 SNPs and cocaine-induced aggression, respectively, were not confounded by sex in Cox regression models (Extended Tables 5 and 6). Not only these associations remained significant when sex was included as a potential confounder in the model, but sex was not associated with the clinical outcomes per se. We also investigated associations between biological sex and both cocaine-induced aggression and transition from 1st cocaine use to CUD using bivariate analyses, showing neither was significant (cocaine-induced aggression mean =1.39 in women and 1.38 in men, median =0 and interquartile range =0-3 in both sexes, Mann-Whitney 4885, p-value = 0.8707; transition from 1st cocaine use to CUD median = 12, interquartile range =0-60 in women and 24 months, interquartile range =0-84 in men, log-rank test Chi2 =1.2, p =0.2).

---

**Extended Data Figure 1.**
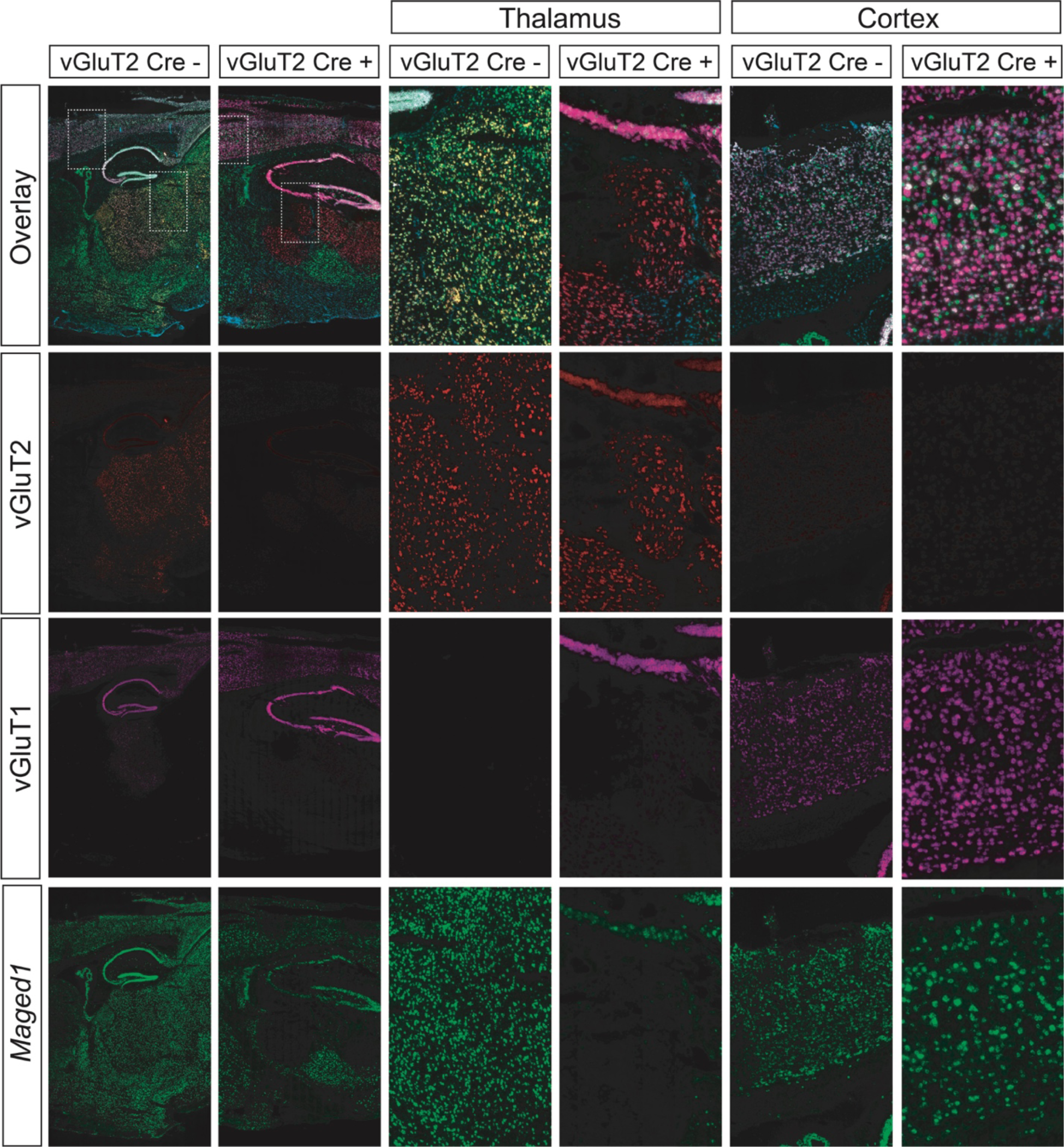
Multiplexed fluorescent in situ hybridization of *Maged1*, vGluT1 and vGluT2. Multiplexed fluorescent in situ hybridization of *Maged1*, vGluT1 and vGluT2 mRNAs. The first column shows a parasagittal slice of a control mouse. The third and fifth columns represent, respectively, a higher magnification focused on the thalamus and on the cortex of the same mouse. We observe that Maged1 is ubiquitously expressed among vGluT1 and vGluT2 cells. It is also expressed in other type of neurons and unmarked cells. The second column shows a *Maged1* KO specifically in vGluT2 expressing neurons. The fourth and sixth columns represent, respectively, a higher magnification focused on the thalamus and on the cortex of the same mouse. We here see an absence of expression of *Maged1* in the thalamic nuclei that is vGluT2 positive, but a normal expression in the other regions, not expressing vGluT2 or expressing vGluT1.

**Extended Data Figure 2.**
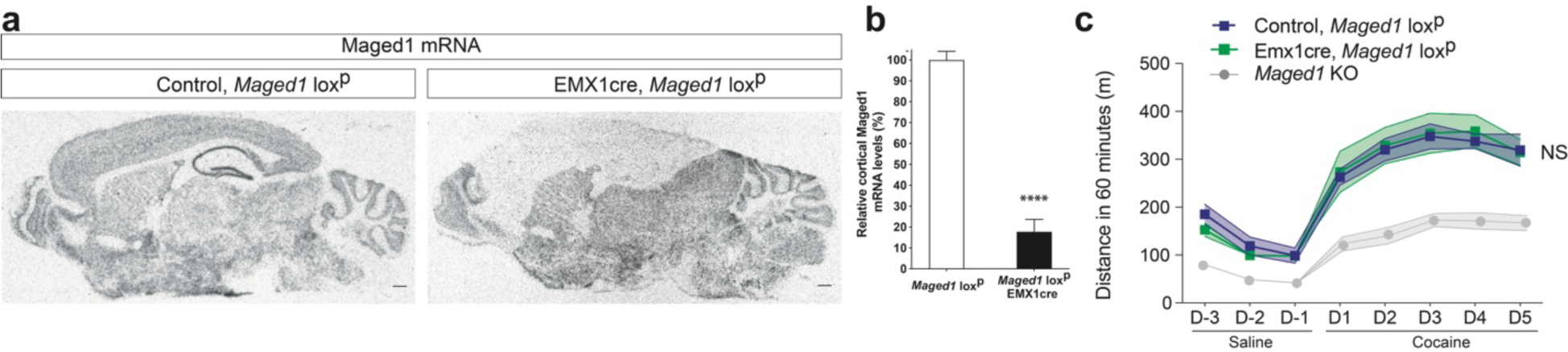
*Maged1* is not necessary for cocaine-induced locomotor sensitization in glutamatergic telencephalic neurons. **a,** In situ hybridization autoradiograms showing *Maged1* mRNA (scale bar: 0.5 mm) **b,** Quantification of the relative cortical optical densities (control, *Maged1*, loxP n = 5; EMX1cre, *Maged1* loxP mice, n = 5; Mann–Whitney test, P < 0.0001) **c,** Cocaine-induced locomotor sensitization (20 mg/kg, ip injection, the mean ± s.e.m., control, *Maged1* loxP, n = 8; EMX1cre, *Maged1* loxP mice, n = 8; repeated measures two-way ANOVA, control versus EMX1cre, *Maged1* loxP, P = 0.9588; days, P<0.0001; interaction factor (genotype X days) P = 0.9519).

**Extended Data Figure 3.**
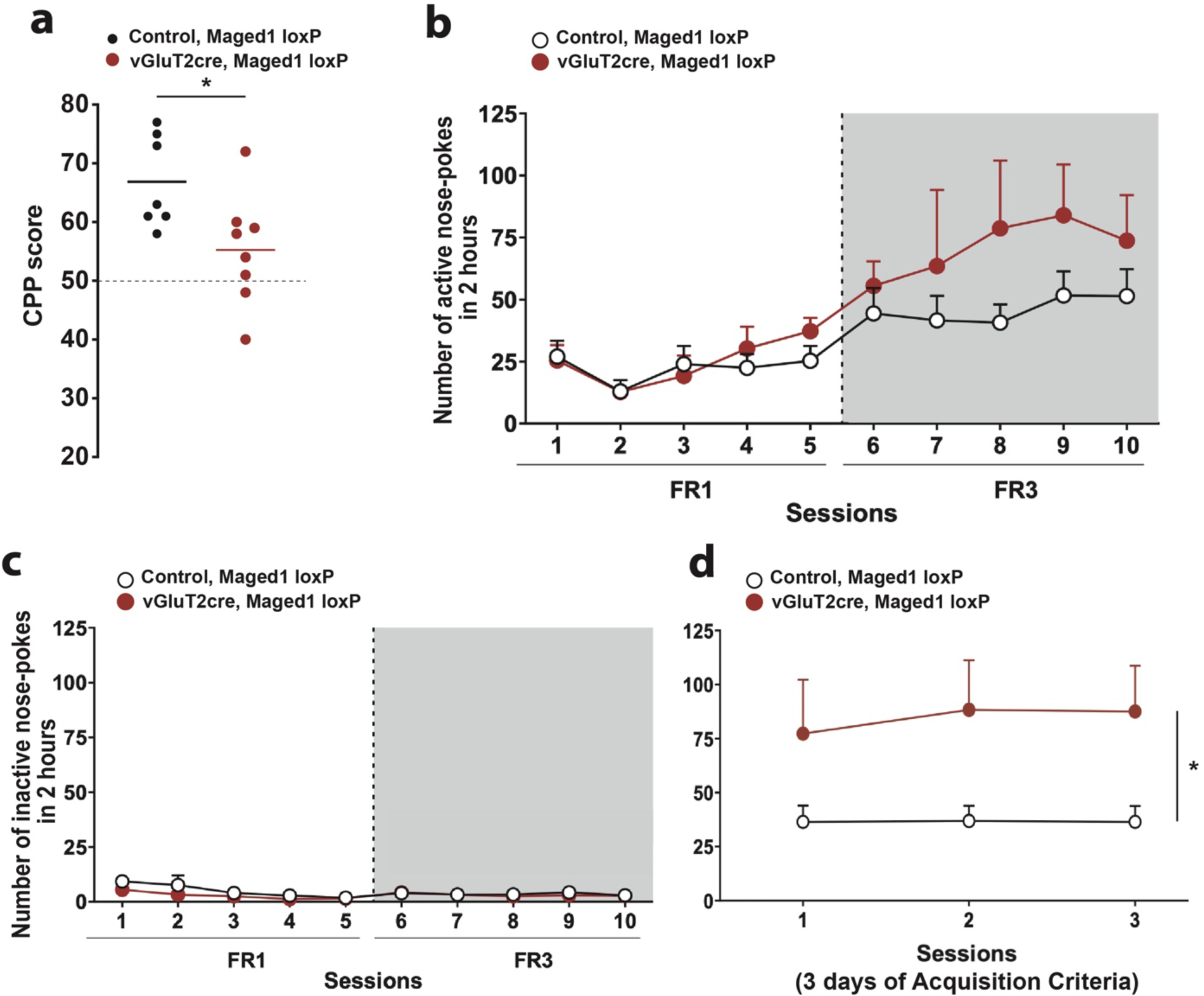
Reinforcing properties of cocaine: cocaine-induced (10mg/kg, ip) conditioned place preference (CPP) and operant conditioning maintained by cocaine (0.5 mg/kg per infusion, iv) self-administration. **a,** CPP scored as the percentage of time spent in the drug-paired compartment (mean ± s.e.m., ncontrol = 8, nvGluT2cre, Maged1loxP = 8, unpaired t-test, *P = 0.0119). **b,** Number of active nose-pokes during the acquisition (fixed ratio 1 (FR1) and 3 (FR3)) of self-administration. Genotype × time F=1.700, n.s., two-way ANOVA repeated measures. **c,** Number of inactive nose-pokes during the acquisition (fixed ratio 1 (FR1) and 3 (FR3)) of self-administration. Genotype × time F=0.2463, n.s., two-way ANOVA repeated measures. **d,** Time course of the active nose-pokes during the three days of accomplishment of acquisition criteria, genotype F=7.746, p<0.05. Data were expressed as mean ± S.E.M; n = 12 control, Maged1 loxp mice and n = 4 vGluT2cre, Maged1 loxP mice.

**Extended Data Figure 4.**
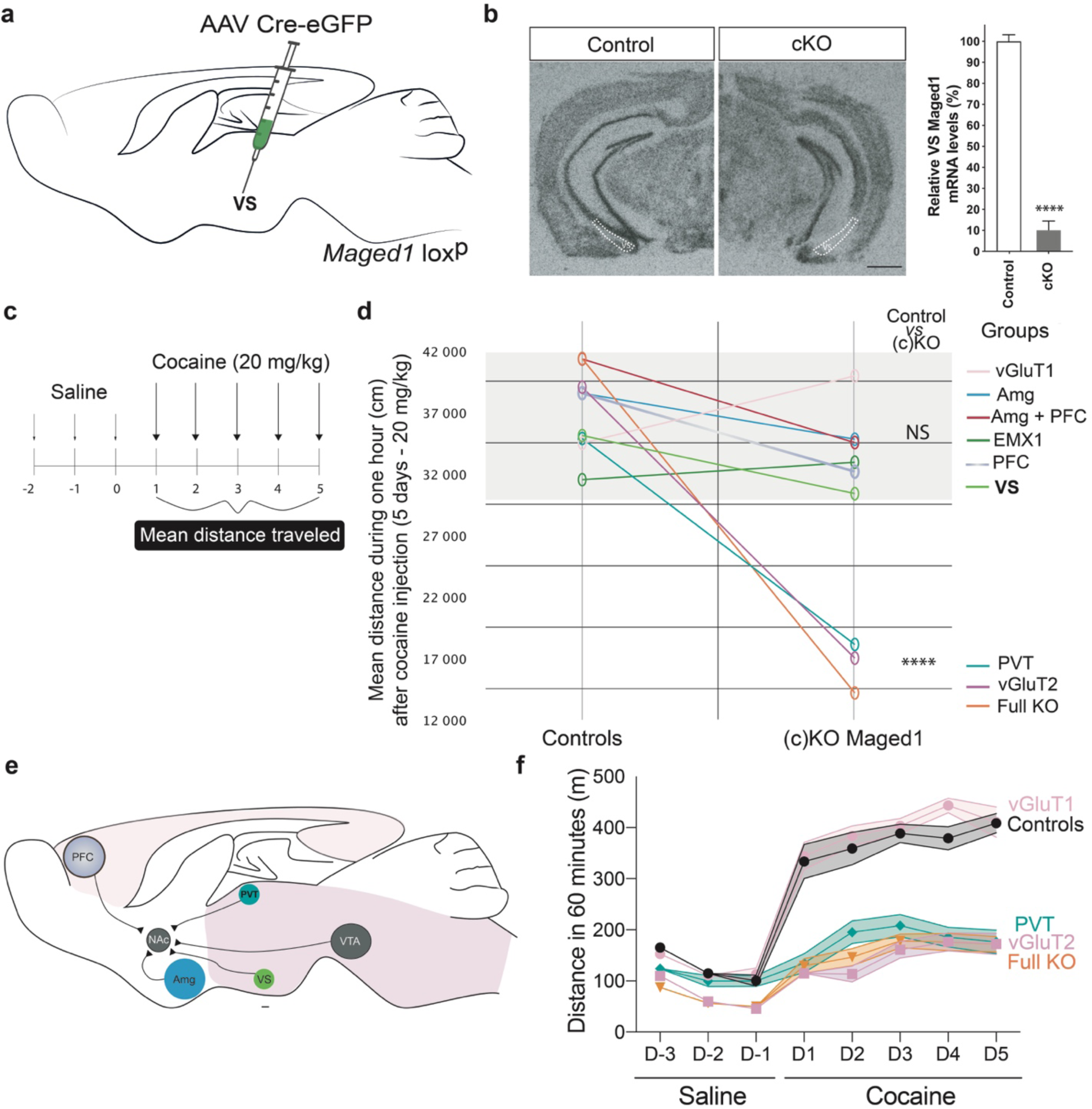
vGluT2 neurons and particularly the paraventricular thalamus are central for *Maged1* role in cocaine-induced behaviors, in contrast with other glutamatergic regions. **a,b,** Ventral subiculum (VS) specific inactivation of *Maged1* with stereotaxic injection of AAV CMV-Cre-eGFP in *Maged1* loxP mice, scheme of the experiment (**a**), in situ hybridization autoradiograms of *Maged1* mRNA (scale bar: 1mm) and quantification of optical densities in the VS (ncontrol = 13, ncKO = 10, P< 0.001, Mann-Whitney U test) (**b**). **c,d,** Summary representation composed by a scheme of the classical cocaine-induced locomotor sensitization protocol, with 3 days of saline injection followed by 5 days of cocaine injection (**c**) and a representation of the mean distance traveled during the 5 days of cocaine injection in the classical sensitization protocol (**d**); segregation between (1) *Maged1* inactivation in the ventral subiculum (VS), the prefrontal cortex (PFC), the cortical and hippocampal EMX1 cells, amygdala (Amg) inactivation, double inactivation in the Amg and PFC, vGluT1 cell inactivation and (2) diencephalic *Maged1* inactivation (PVT and vGluT2) (two-way ANOVA, groups, P < 0.0001; treatment, P < 0.0001; interaction factor, P < 0.0001; Sidak’s posttest, ****P < 0.0001). **e,** Color scheme of a sagittal section of the mouse brain that represents the different regions with targeted *Maged1* inactivation. **f,** Superimposed cocaine-induced locomotor sensitization experiment showing clear superimposition of full *Maged1*-KO mice with the vGluT2 mice and of the vGluT1 mice with the control mice.

**Extended Data Figure 5.**
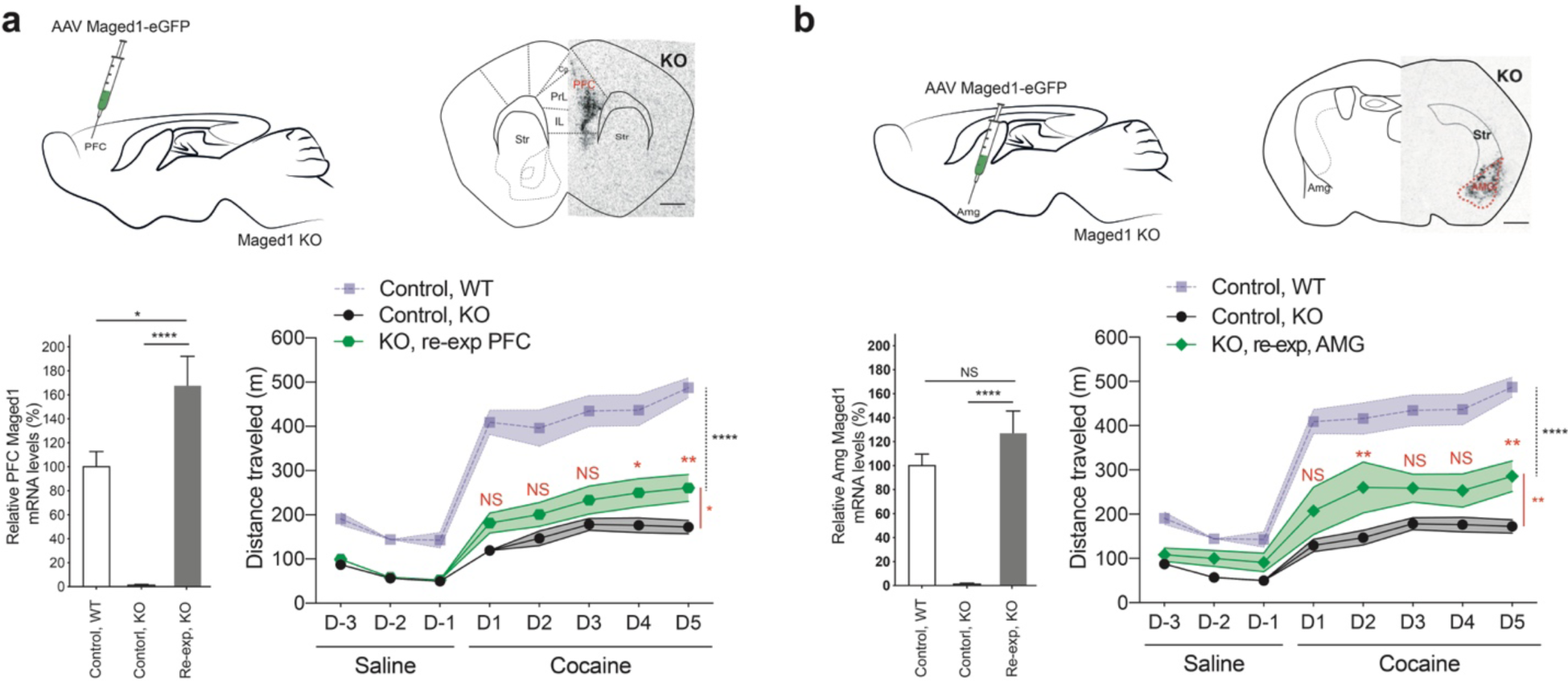
Partial effect of *Maged1* re-expression in glutamatergic vGluT1/2 nuclei. **a,** PFC specific re-expression (re-exp) of *Maged1* with stereotaxic injection of AAV CMV-*Maged1*-eGFP in *Maged1* KO mice, scheme of the experiment. In situ hybridization autoradiograms of *Maged1* mRNA (scale bar: 1mm) and quantification of optical densities in the PFC (ncontrolWT = 9, ncontrolKO = 11, nre-expKO = 16 P< 0.0001, Kruskal-Wallis test). Cocaine induced locomotor sensitization (20 mg/kg, ip, mean ± s.e.m., control WT n = 9, control KO mice n = 15, KO, PFC, AAV-eGFP-*Maged1* = 16, mixed-effects model, control KO versus re-expression KO P = 0.0268, days P<0.0001, interaction factor (genotype X days) P = 0.0046, Sidak’s post-test *P < 0.05 **P < 0.01). **b,** Amg specific re-exp of *Maged1* with stereotaxic injection of AAV CMV-*Maged1*-eGFP in *Maged1* KO mice, scheme of the experiment. In situ hybridization autoradiograms of *Maged1* mRNA (scale bar: 1mm) and quantification of optical densities in the Amg (ncontrolWT = 9, ncontrolKO = 15, nre-expKO = 5 P< 0.0001, Kruskal-Wallis test). Cocaine induced locomotor sensitization (20 mg/kg, ip, mean ± s.e.m., control WT n = 9, control KO mice n = 15, KO, Amg, AAV-eGFP-*Maged1* = 5, mixed-effects model, control KO versus re-exp KO P = 0.0020, days P<0.0001, interaction factor (genotype X days) P = 0.0733, Sidak’s post-test **P < 0.01).

**Extended Data Figure 6.**
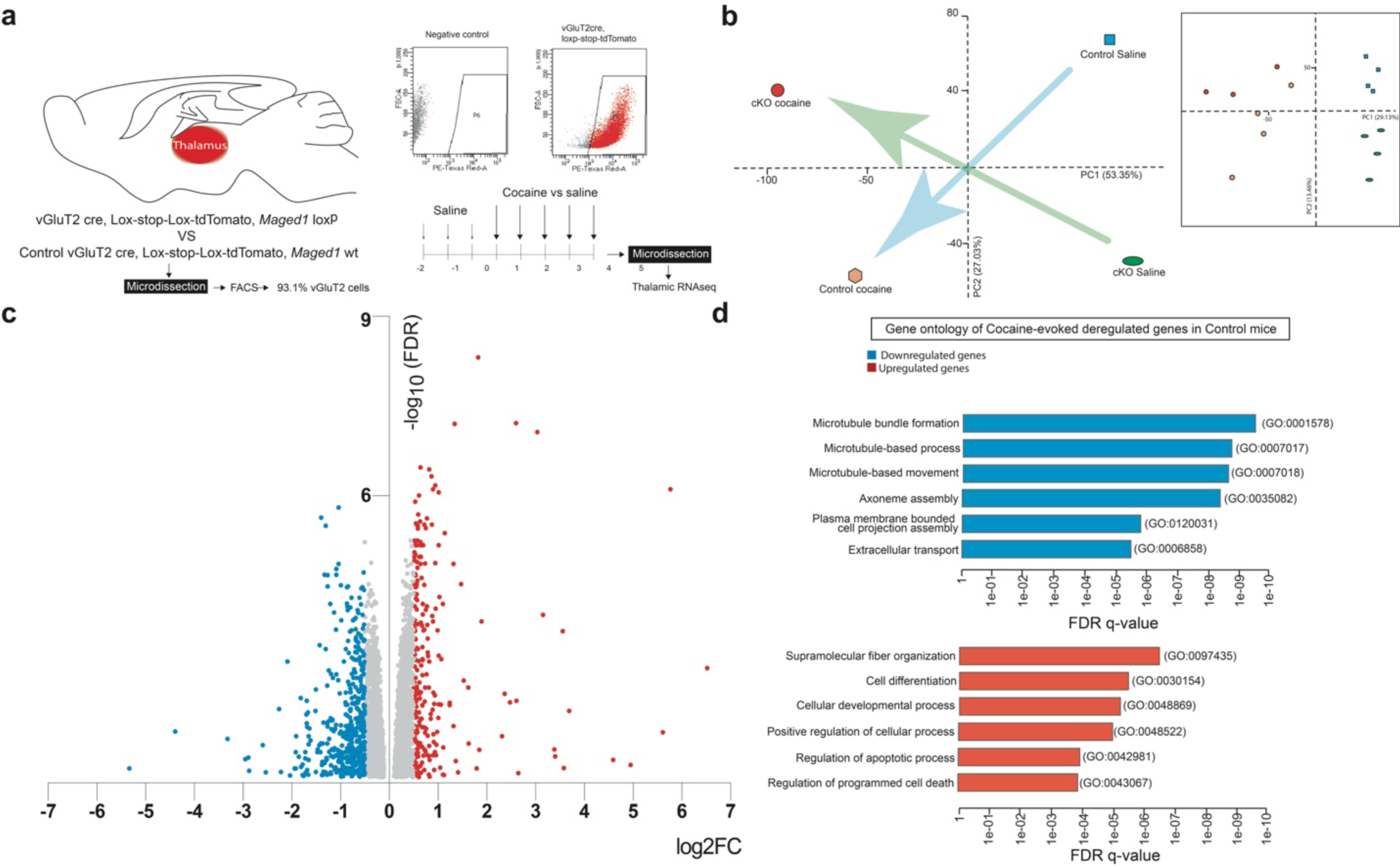
Flow cytometry of the thalamus followed by transcriptomic and gene ontology analysis. **a,** Scheme of the RNA-seq experiment. Flow cytometry showed that 93.1% of microdissected thalamic tissue consisted of vGluT2 cells (n = 6). **b,** Principal component analysis revealed four clusters referring to control and vGluT2cre, *Maged1* loxP (cKO) treated either with saline or with cocaine during 5 consecutive days. **c,** Volcano plot showing the significantly differentially expressed genes between control saline and control cocaine mice. (452 downregulated and 252 upregulated genes, log2|fold change, FC|≥0.5, false discovery rate (FDR)<0.05). **d,** Gene Ontology analysis (biological process) showing categories with statistically significant enrichment of downregulated (top) and upregulated (bottom) genes following cocaine administration in control mice.

**Extended Data Figure 7.**
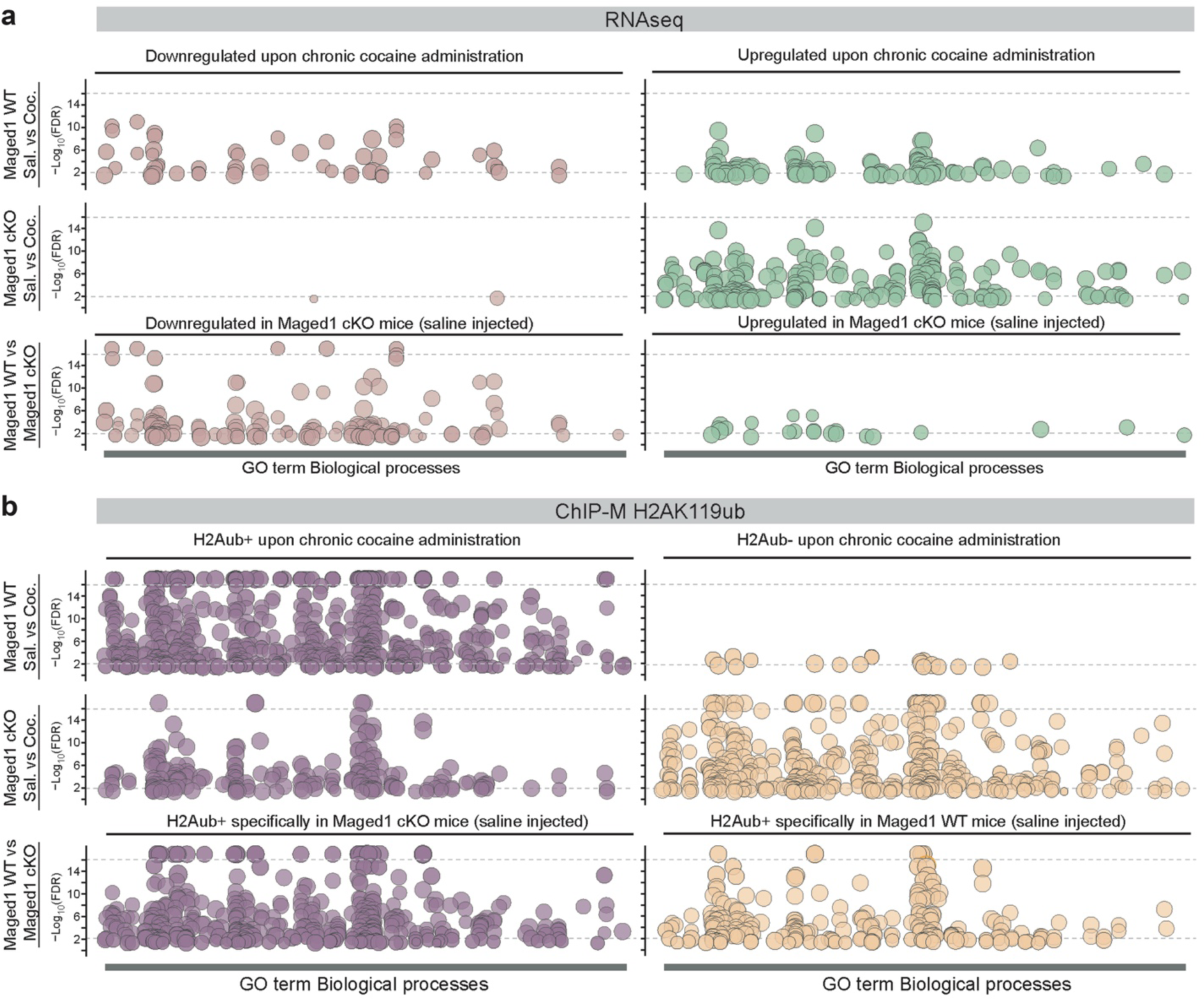
Gene ontology terms – an overview. Graphical overview of significant gene ontology (GO) terms and their –Log10(false discovery rate, FDR) found in all the comparisons of conditions (saline versus chronic cocaine in control mice and *Maged1* cKO and control versus *Maged1* cKO) associated to significant transcriptomic modifications (RNAseq, **a**) and H2Aub enrichment (ChIP-M H2AK119ub, **b**). The GO terms are organized on the X-axis by their identifier number.

**Extended Data Figure 8.**
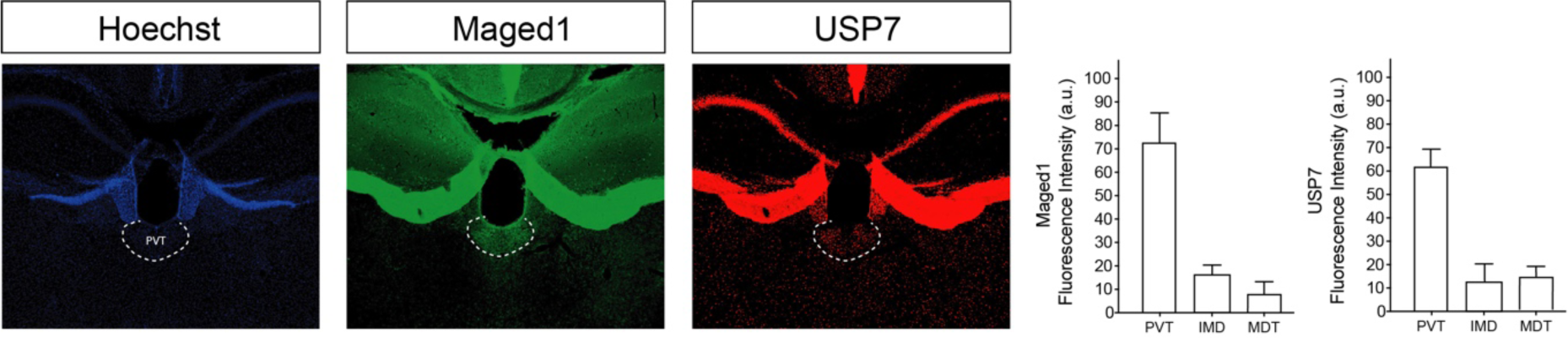
Relatively high immunohistochemistry signal of Maged1 and USP7 in the PVT. Immunohistochemistry staining of Maged1 (green) and USP7 (red) (Hoechst staining (blue)) showing relative increases in USP7 and Maged1 signals in the PVT. For quantification, we used n = 5 mice (for Maged1: Repeated-measure one-way ANOVA, ***p = 0.002, Tukey’s multiple comparisons test, PVT vs IMD, **P = 0.002; PVT vs MDT, **P = 0.0027; IMD vs MDT P = 0.1552; for USP7: Repeated-measure one-way ANOVA, ****p < 0.0001, Tukey’s multiple comparisons test, PVT vs IMD, **P = 0.0013; PVT vs MDT, ***P = 0.0006; IMD vs MDT P = 0.7814). PVT: the paraventricular nucleus of the thalamus, MDT: the mediodorsal thalamic nucleus (including the medial, lateral and central parts), IMD: the intermediodorsal nucleus of thalamus.

**Extended Data Figure 9.**
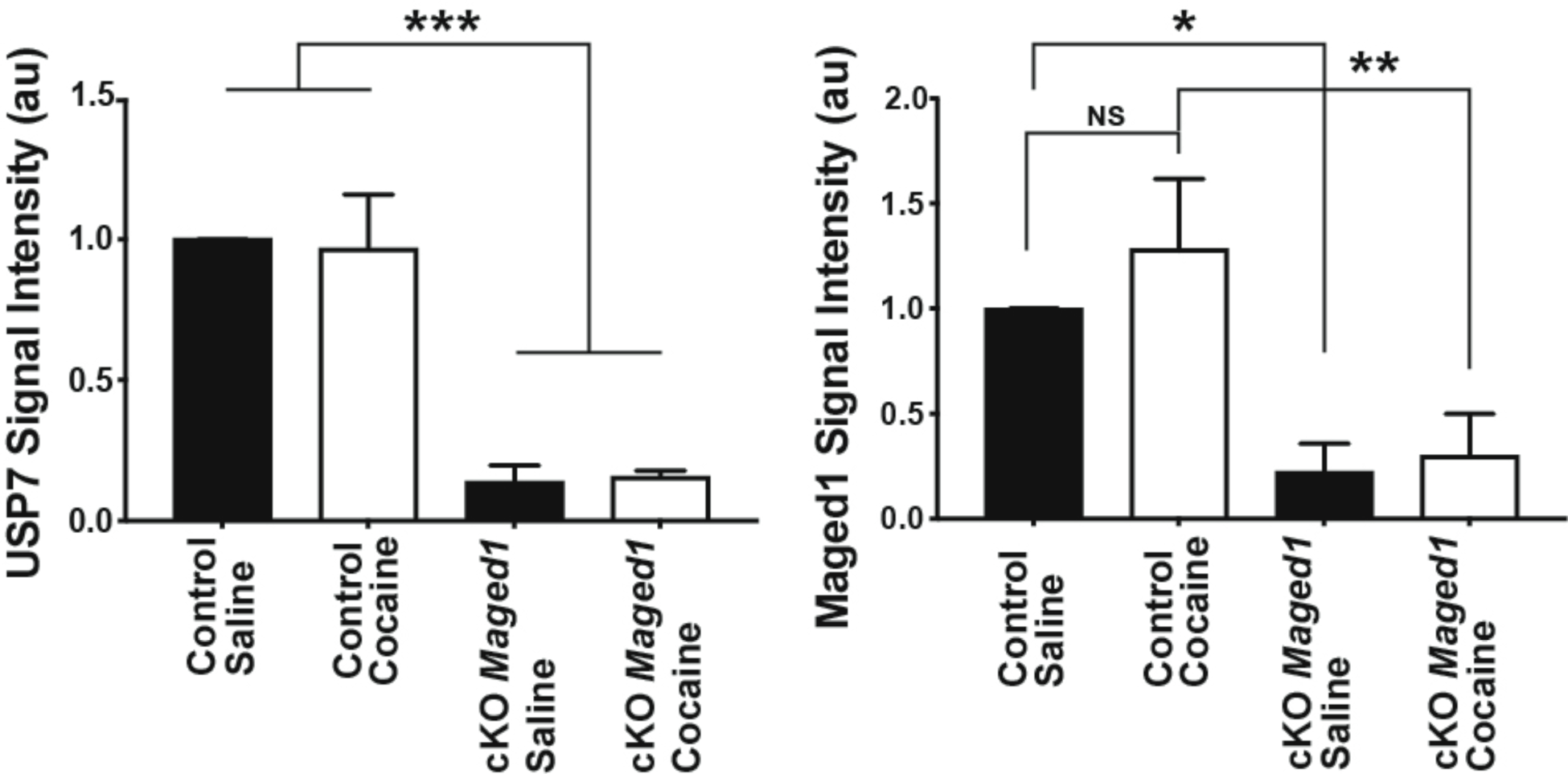
Immunoprecipitation of H2Aub, western blotting quantification. MnIP of monoubiquitinated H2AK119 (H2Aub) and its partners with identification of Maged1 and USP7 by western blot. Quantification of USP7 and Maged1 signal intensities relative to the control condition (mean ± s.e.m., n = 5) (one-way ANOVA, P = 0.001 and Sidak’s post-test *P < 0.05 **P < 0.01 ***P < 0.001).

**Extended Data Figure 10.**
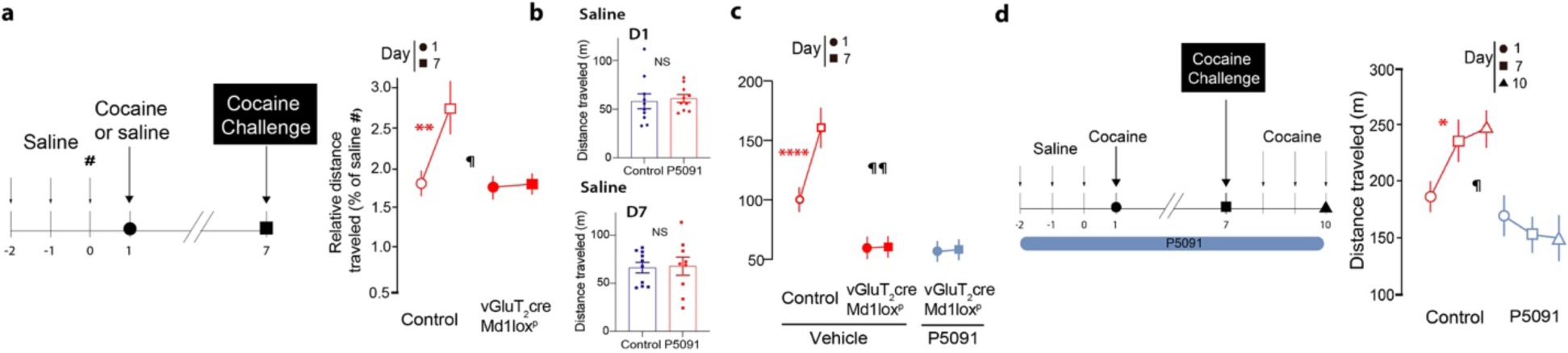
Long-lasting abolition of cocaine-induced sensitization induced by USP7 inhibition, similarly to *Maged1* conditional KO, with no effect on spontaneous locomotor activity. **a,** Scheme of cocaine sensitization (cocaine, intraperitoneal (ip) injection, 15 mg/kg, without P5091). Cocaine-induced sensitization normalized on saline day 0 (#) was abolished after inactivation of *Maged1* in vGluT2 neurons (cKO, vGluT2cre *Maged1* loxp) (the mean ± s.e.m., control, n = 9, cKO, n = 8; repeated measures two-way ANOVA, control versus cKO, P = 0.1029; days, P < 0.0301; interaction factor (genotype X days) ¶P = 0.0370, Sidak’s posttest (D1 versus D7) **P = 0.0077). **b,** Locomotor activity (m) during 30 minutes after saline injection after 4 (top) and 10 (bottom) daily P5091 systemic administration, showing that P5091 itself had no effect on spontaneous locomotor activity during the total length of the protocol. **c,** Cocaine-induced sensitization is abolished after Maged1 inactivation in vGluT2 cells (vGluT2cre Maged1 loxp) (vehicle and P5091 administration, ip, 10 mg/kg (mean ± s.e.m., ncontrol = 9, nvGluT2cre *Maged1* loxp, vehicle = 8, vGluT2cre *Maged1* loxp, P5091 = 8, repeated measures two-way ANOVA, group factor, P < 0.0001; days P = 0.0052; interaction factor (group X days) ¶¶P = 0.0020, Sidak’s post-test (J1 versus J7) NS for vGluT2cre Maged1 loxp, vehicle and P5091, ****P<0.0001 for control and vehicle). **d,** Scheme of the cocaine sensitization (cocaine, intraperitoneal (ip), 15 mg/kg) during USP7 inhibition experiment (P5091, ip, 10 mg/kg). Cocaine-induced sensitization is abolished after systemic inhibition of USP7 (mean ± s.e.m., ncontrol = 10, nP5091 = 8, repeated measures two-way ANOVA, vehicle versus P5091, P < 0.0001; days P = 0.4444; interaction factor (P5091 X days) ¶P = 0.0495, Sidak’s post-test (J1 versus J10), * P = 0.0288).

**Extended Data Figure 11.**
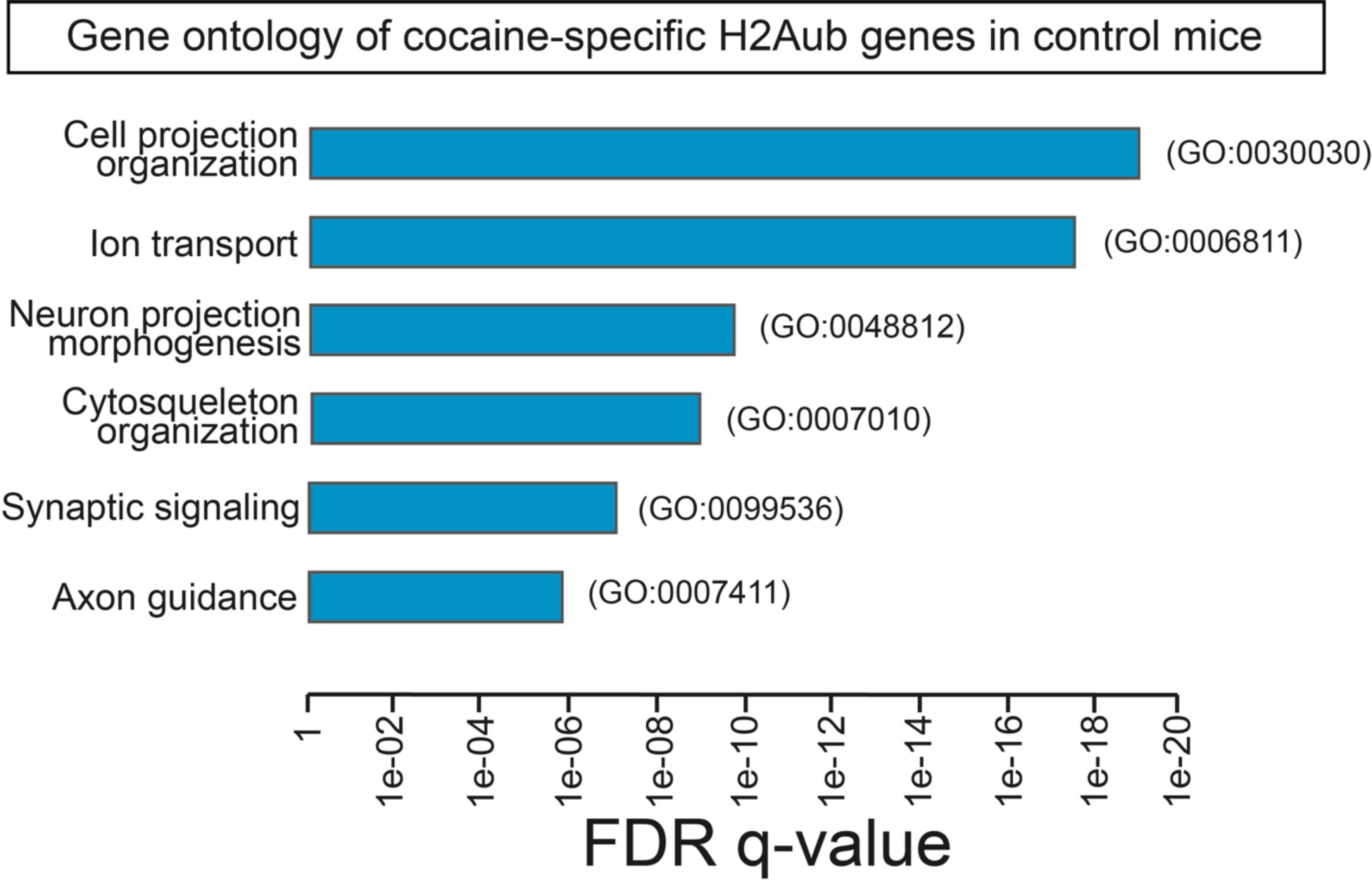
Gene ontology analysis of H2Aub enrichment. Bar plot presenting the gene ontology terms (biological process) associated with genes with H2Aub enrichment specifically in *Maged1* control mice injected with cocaine.

**Extended Data Figure 12.**
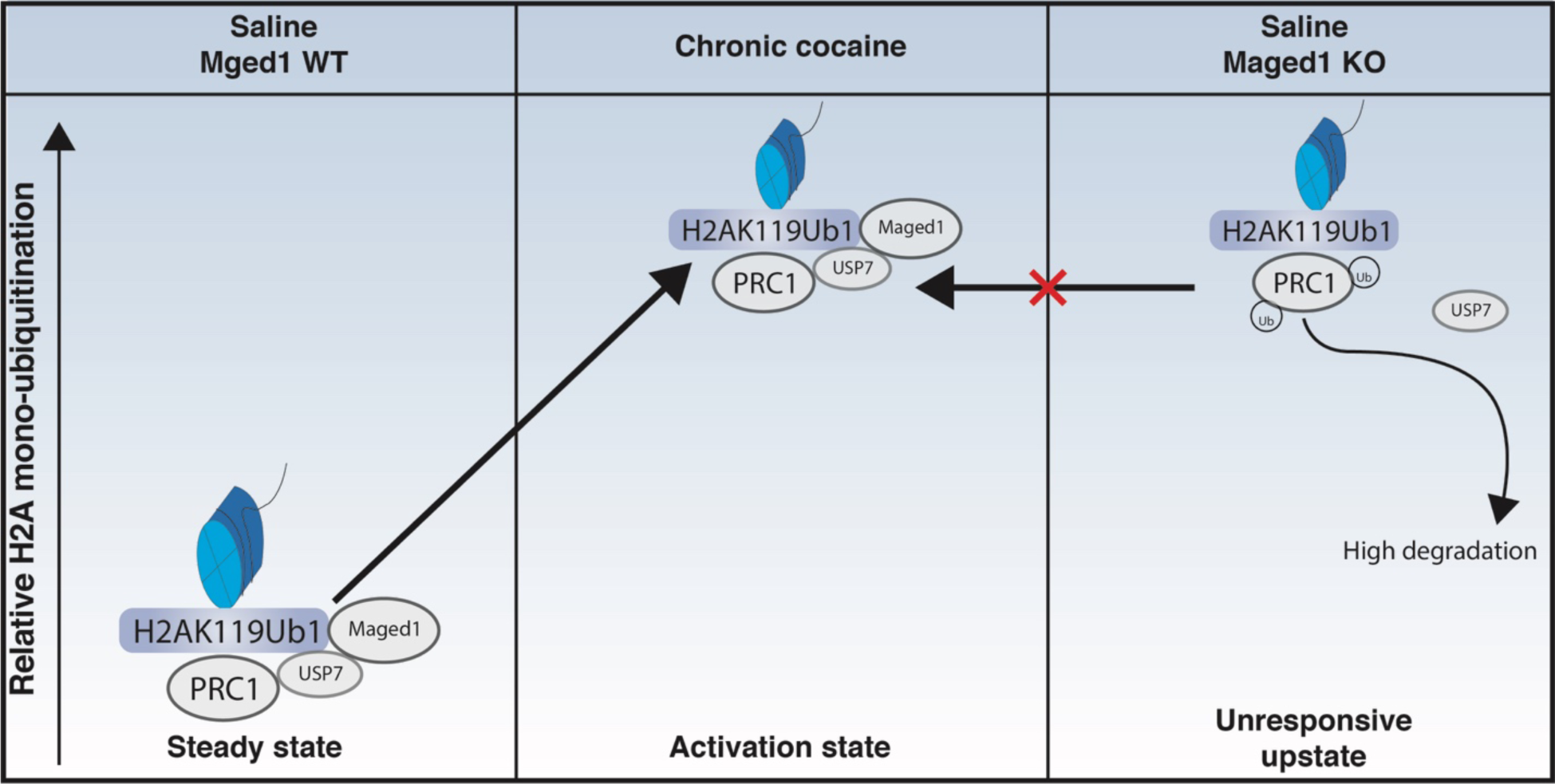
Proposed mechanistic model for the role of Maged1 and USP7 in cocaine use disorder. In this model, polycomb repressive complex 1 (PRC1), with Maged1 and USP7 are needed for cocaine-induced H2A mono-ubiquitination (H2AK119ub1). Maged1 enables the interaction between USP7 and H2AK119ub1, possibly controlling the fate of PRC1. Our results suggest that Maged1 helps, together with USP7, maintaining PRC1 in a responsive native steady state. When cocaine is administered, the complex is activated, globally leading to H2A mono-ubiquitination. However, in the absence of Maged1, USP7 cannot maintain PRC1 in a responsive steady sate and the complex is blocked in an unresponsive upstate where upstream events (like cocaine exposure) do not modify its state.

**Extended Data Figure 13.**
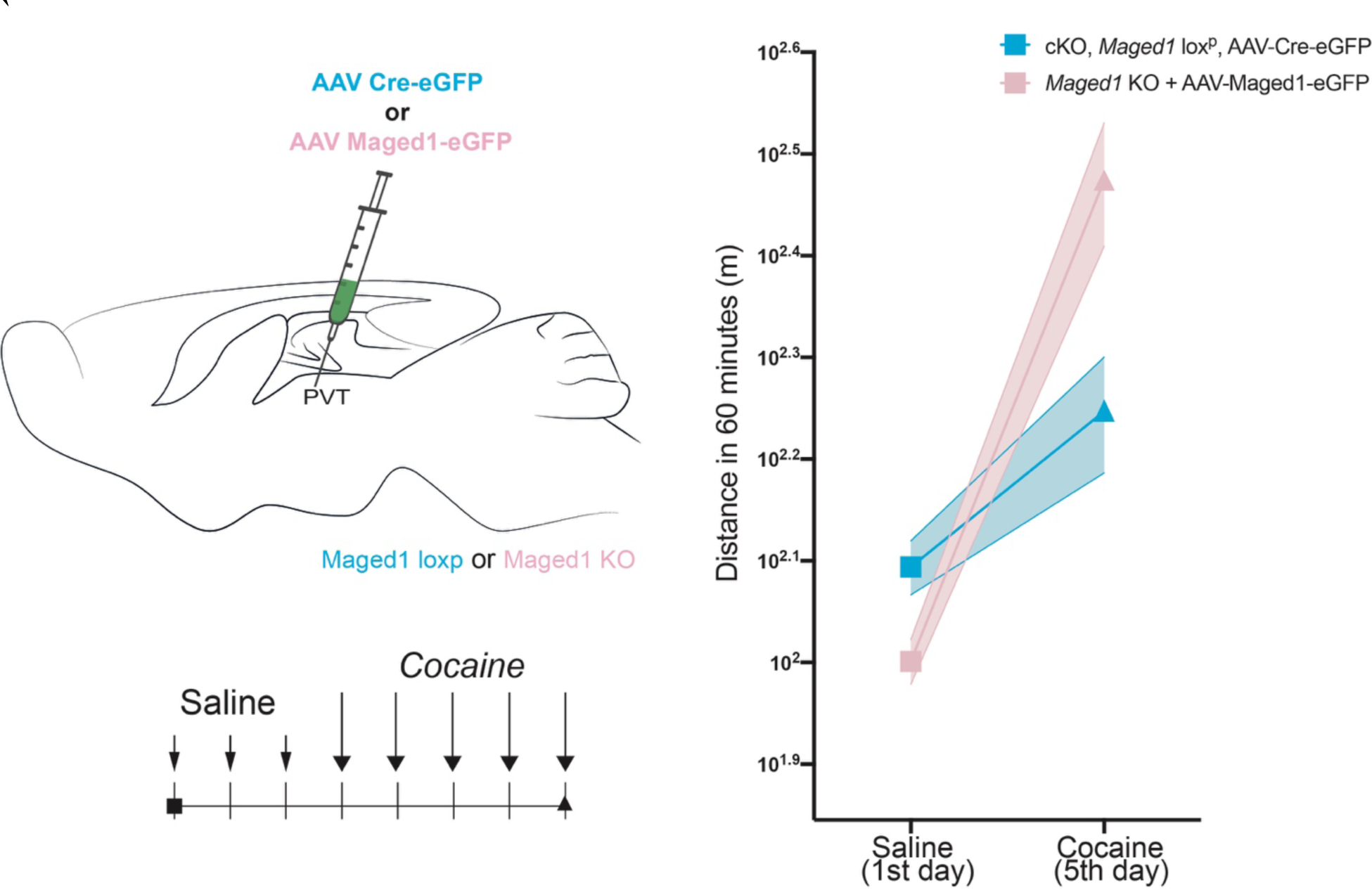
Cocaine-induced locomotor sensitization is modulated by Maged1 in the thalamus (PVT). Comparison of the locomotor activity on the first day of saline injection and the last day of cocaine injection in the rescue model and in the Maged1 cKO model (two-way ANOVA, Maged1 KO, AAV-Maged1 (PVT) versus Maged1 loxP, AAV-Cre (PVT), P = 0.0552; days, P<0.0001; interaction factor (genotype X days) P = 0.0037)

**Extended Data Figure 14.**
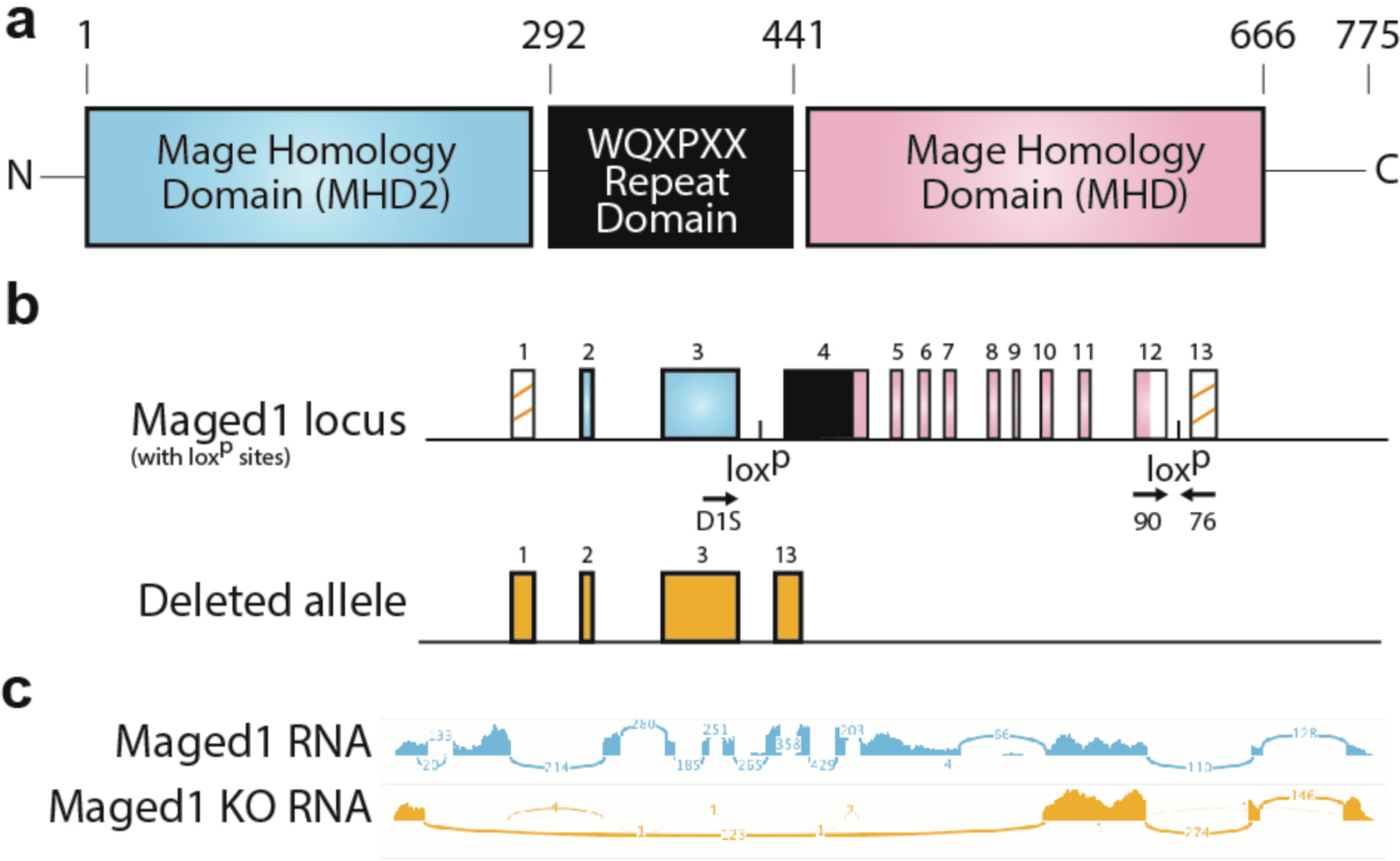
Maged1 domains with their respective mRNA and DNA. **a,** The Maged1 protein is composed of the conserved Mage Homology Domains (MHD) and another MHD, MHD2 (poorly conserved between species and common to some Maged family members with no significant homology to other proteins) separated by a hexapeptide repeated domain. **b,** Schematic representation of the Maged1 locus and the knock-out allele. Boxes represent exons. The spliced part of the locus is represented with orange, with the MHD in black. Two LoxP sites were inserted into the Maged1 locus, the first one between the exons 3 and 4 and the second one between the exons 12 and 13. After recombination by a Cre recombinase, the exons 4 to 12 are deleted to obtain the (functional) Maged1 knock-out (KO) allele. Arrows represent PCR primers used for genotyping. In this work, primers 90 and 76 were used to detect wild-type and conditioned (loxp) alleles whereas the couple D1S and 76 detect the KO allele^50^. **c,** RNAseq Mapping on Maged1 gene for control and KO mice (after FACS sorting).

**Extended Data Figure 15.**
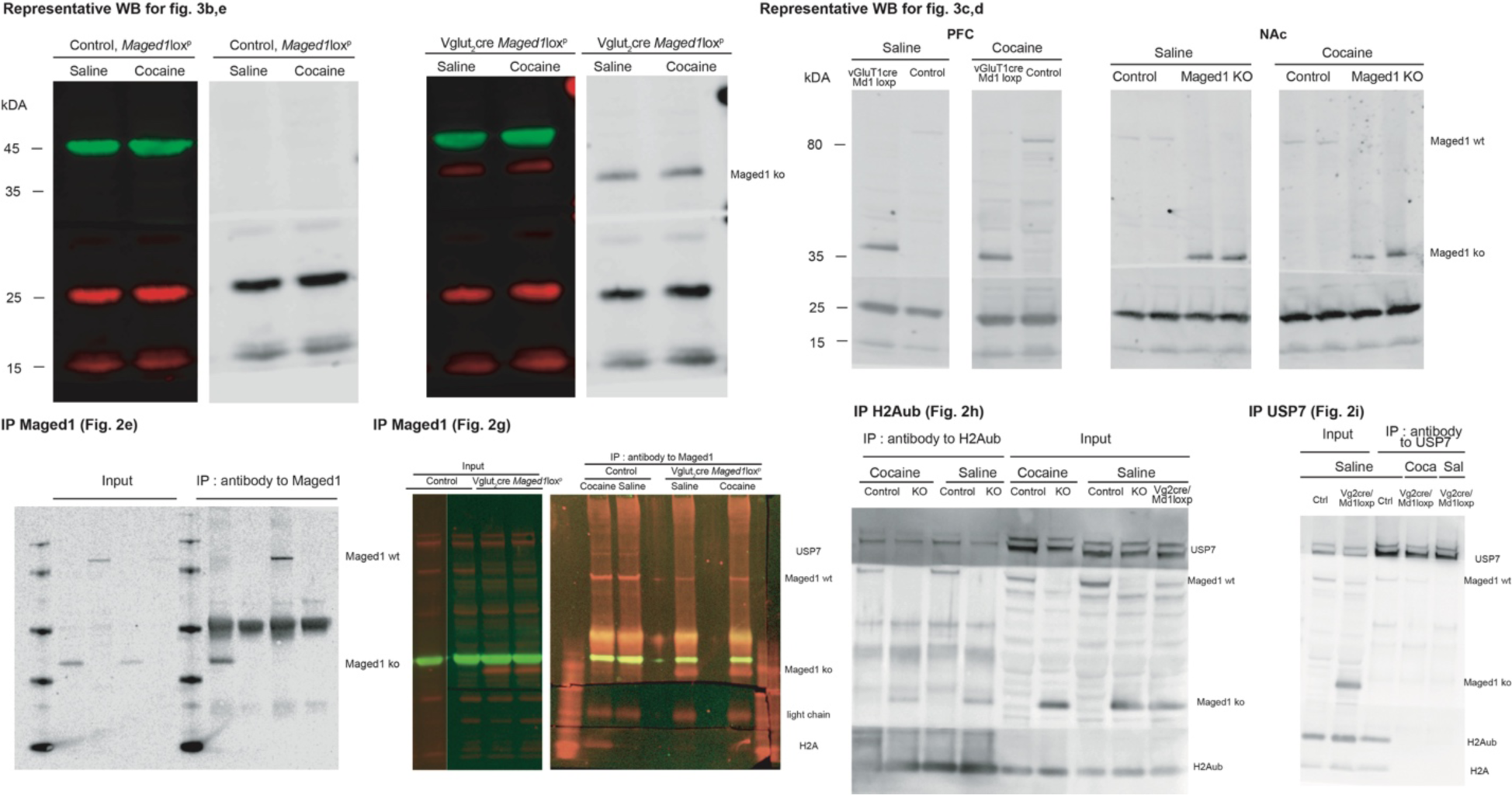
Raw immunoblots of immunoprecipitations shown in. Fig. 2 **and representative western blot analyzed in** Fig. 3.

**Extended Data Fig. 16.**
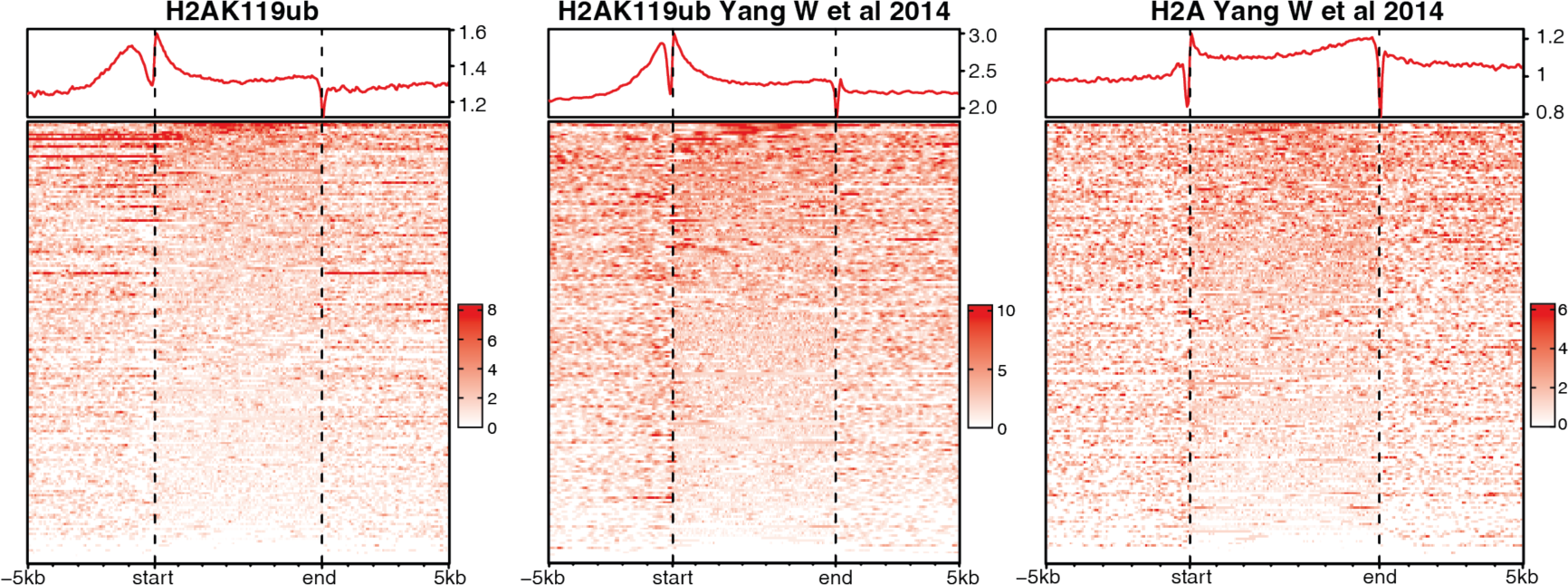
Coverage heatmap plots comparing the genome wide distribution of H2Aub over gene bodies and upstream and downstream regions reported in our study (left) with those of H2Aub and H2A form datasets of Yang W et al., 2014 (center and right).

**Extended Data Table 1.** RNAseq and ChIPseq gene lists, Gene ontology (GOterm) and curated gene sets.

*Please refer to extensive lists in “TableS1.xlsx” file*

**Extended Data Table 2.** Mass spectrometry results (mice and humans) from Maged1/MAGED1 IP, Maged1/MAGED1 partners protein lists.

*Please refer to extensive lists in “TableS2.xlsx” file*

**Extended Data Table 3.** Description of the clinical sample. *All SUD diagnoses are lifetime and correspond to DSM-IV TR dependence. SAPS, Scale for Assessment of Positive Symptoms – Cocaine-Induced Psychosis; IQR, interquartile range.*

|  | N (%) or mean (SD) | N |
| --- | --- | --- |
| Sex | 82 (23%) | 350 |
| Age | 38 (9) | 350 |
| Age at first cocaine use | 23 (7) | 338 |
| Age at onset of cocaine use disorder | 27 (8) | 261 |
| Number of cocaine uses during first year after initiation | 25 (82) | 203 |
| Transition between first cocaine use and cocaine use disorder (months) | 52 (72) | 260 |
| SAPS score – aggression | 1.4 (1.7) | 240 |
| SAPS score – physical symptoms before cocaine use | 1.3 (1.5) | 240 |
| Current tobacco smoking | 304 (87%) | 350 |
| Alcohol use disorder | 210 (62%) | 338 |
| Opioid use disorder | 208 (61%) | 340 |
| Cannabis use disorder | 217 (64%) | 338 |
| Benzodiazepine use disorder | 140 (41%) | 339 |
| Number of comorbid substance use disorders (tobacco excluded) | 1: 50 (14%) | 350 |
|  | 2: 55 (16%) |  |
|  | 3: 89 (26%) |  |
|  | 4: 91 (26%) |  |
|  | 5: 64 (18%) |  |
| Number of psychotropic medications in current treatment (median, IQR) | 1 (0-1) | 350 |
| Number of non-psychotropic medications in current treatment (median, IQR) | 1 (0-2) | 350 |

**Extended Data Table 4.**
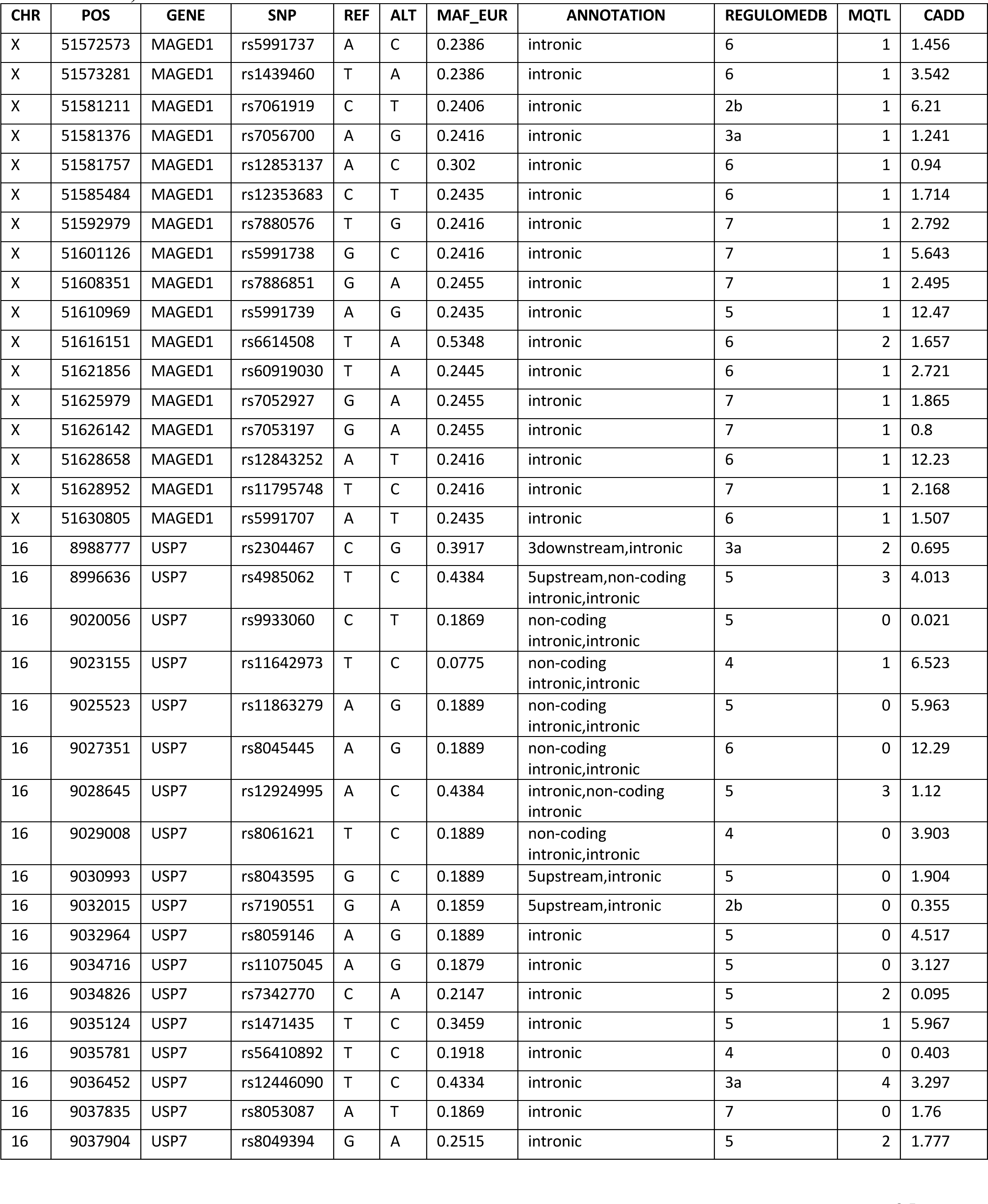
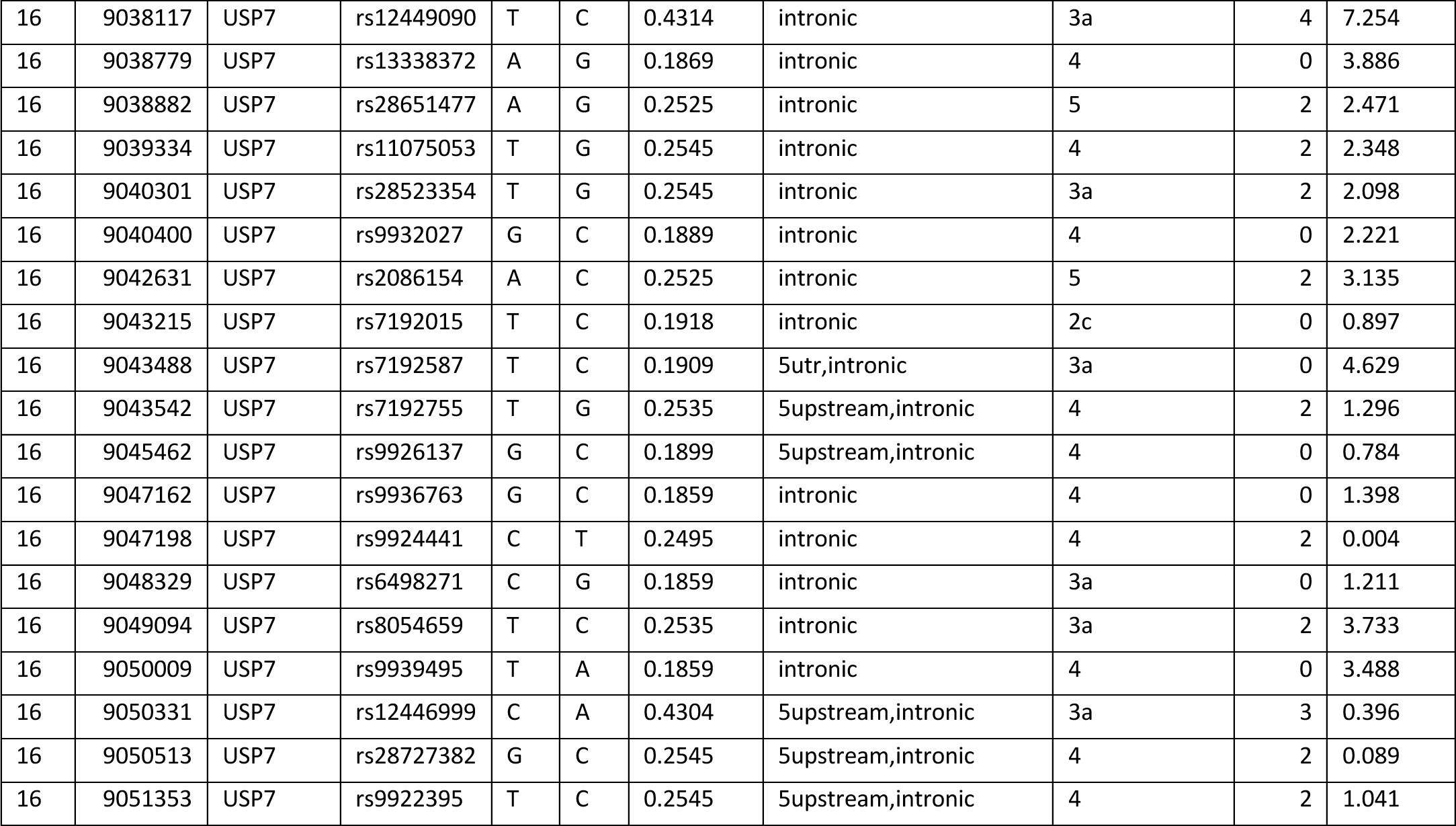
Functional analysis of 54 SNPs showing significant associations (at FDR <0.05)

**Extended Data Table 5.** Cox regression with the transition from first cocaine use to cocaine use disorder (CUD) as the dependent variable and *MAGED1* rs7886851-A as the lead SNP. Hazard ratios (HR) and 95% confidence intervals (CI) are displayed. N =127.

| Characteristic | HR | 95% CI | p-value |
| --- | --- | --- | --- |
| rs7886851 | 1.52 | 1.06, 2.19 | 0.024 |
| Age | 0.96 | 0.94, 0.98 | <0.001 |
| Sex | 0.82 | 0.55, 1.23 | 0.3 |
| Impulsive temper | 1.00 | 1.00, 1.01 | 0.4 |

**Extended Data Table 6.** Cox regression with the transition from first cocaine use to cocaine use disorder (CUD) as the dependent variable and *MAGED1* rs6614508-A as the lead SNP. Hazard ratios (HR) and 95% confidence intervals (CI) are displayed. N =127.

| **Characteristic** | **HR** | **95% CI** | **p-value** |
| --- | --- | --- | --- |
| rs6614508 | 0.56 | 0.36, 0.86 | 0.007 |
| Age | 0.95 | 0.92, 0.98 | <0.001 |
| Sex | 0.90 | 0.57, 1.43 | 0.7 |
| Impulsive temper | 1.00 | 0.99, 1.01 | 0.5 |

**Extended Data Table 7.** Gene expression in the brain and SNPs from *MAGED1* and *USP7* with significant associations with cocaine-related phenotypes according to http://www.braineac.org/.

*Please refer to extensive lists in “TableS7.xlsx” file*

**Extended Data Table 8.** Summary statistics of associations between *MAGED1* and *USP7* SNPs and seven cocaine-related phenotypes.

*Please refer to extensive lists in “Table8.xlsx” file*

## References

1. Nestler, E. J. & Lüscher, C. The Molecular Basis of Drug Addiction: Linking Epigenetic to Synaptic and Circuit Mechanisms. Neuron 102, 48–59 (2019).

2. Hamilton, P. J. & Nestler, E. J. Epigenetics and addiction. Current Opinion in Neurobiology 59, 128–136 (2019).

3. Maze, I. et al. Cocaine dynamically regulates heterochromatin and repetitive element unsilencing in nucleus accumbens. PNAS 108, 3035–3040 (2011).

4. Hamilton, P. J. & Nestler, E. J. Epigenetics and addiction. Current Opinion in Neurobiology 59, 128–136 (2019).

5. Cheron, J. & Kerchove d’Exaerde, A. de. Drug addiction: from bench to bedside. Transl Psychiatry 11, 1–14 (2021).

6. Xu, S.-J. et al. Chromatin-mediated alternative splicing regulates cocaine-reward behavior. Neuron 109, 2943–2966.e8 (2021).

7. Lepack, A. E. et al. Dopaminylation of histone H3 in ventral tegmental area regulates cocaine seeking. Science 368, 197–201 (2020).

8. Renthal, W. et al. Genome-wide analysis of chromatin regulation by cocaine reveals a role for sirtuins. Neuron 62, 335–348 (2009).

9. De Backer, J.-F. et al. Deletion of Maged1 in mice abolishes locomotor and reinforcing effects of cocaine. EMBO Rep. 19, (2018).

10. Kim, R. Q. & Sixma, T. K. Regulation of USP7: A High Incidence of E3 Complexes. J Mol Biol 429, 3395–3408 (2017).

11. Kampman, K. M. The treatment of cocaine use disorder. Science Advances 5, eaax1532.

12. Gee, R. R. F. et al. Emerging roles of the MAGE protein family in stress response pathways. Journal of Biological Chemistry 295, 16121–16155 (2020).

13. Doyle, J. M., Gao, J., Wang, J., Yang, M. & Potts, P. R. MAGE-RING protein complexes comprise a family of E3 ubiquitin ligases. Mol Cell 39, 963–974 (2010).

14. Robinson, T. E. & Berridge, K. C. The neural basis of drug craving: an incentive-sensitization theory of addiction. Brain Res Brain Res Rev 18, 247–291 (1993).

15. Valjent, E. et al. Mechanisms of locomotor sensitization to drugs of abuse in a two-injection protocol. Neuropsychopharmacology 35, 401–415 (2010).

16. Creed, M., Pascoli, V. J. & Lüscher, C. Refining deep brain stimulation to emulate optogenetic treatment of synaptic pathology. Science 347, 659–664 (2015).

17. Schenk, S. & Partridge, B. Sensitization and tolerance in psychostimulant self-administration. Pharmacol Biochem Behav 57, 543–550 (1997).

18. Vezina, P., Lorrain, D. S., Arnold, G. M., Austin, J. D. & Suto, N. Sensitization of midbrain dopamine neuron reactivity promotes the pursuit of amphetamine. J Neurosci 22, 4654–4662 (2002).

19. De Vries, T. J., Schoffelmeer, A. N., Binnekade, R., Mulder, A. H. & Vanderschuren, L. J. Drug-induced reinstatement of heroin- and cocaine-seeking behaviour following long-term extinction is associated with expression of behavioural sensitization. Eur J Neurosci 10, 3565–3571 (1998).

20. Suto, N. et al. Previous exposure to psychostimulants enhances the reinstatement of cocaine seeking by nucleus accumbens AMPA. Neuropsychopharmacology 29, 2149–2159 (2004).

21. Chisholm, A. et al. Assessing the Role of Corticothalamic and Thalamo-Accumbens Projections in the Augmentation of Heroin Seeking in Chronically Food-Restricted Rats. J Neurosci 41, 354–365 (2021).

22. Otis, J. M. et al. Paraventricular Thalamus Projection Neurons Integrate Cortical and Hypothalamic Signals for Cue-Reward Processing. Neuron 103, 423–431.e4 (2019).

23. Penzo, M. A. & Gao, C. The paraventricular nucleus of the thalamus: an integrative node underlying homeostatic behavior. Trends in Neurosciences (2021) doi:10.1016/j.tins.2021.03.001.

24. Perez, S. M. & Lodge, D. J. Convergent Inputs from the Hippocampus and Thalamus to the Nucleus Accumbens Regulate Dopamine Neuron Activity. J. Neurosci. 38, 10607–10618 (2018).

25. Everitt, B. J. & Robbins, T. W. Drug Addiction: Updating Actions to Habits to Compulsions Ten Years On. Annual Review of Psychology 67, (2015).

26. Schuettengruber, B., Bourbon, H.-M., Croce, L. D. & Cavalli, G. Genome Regulation by Polycomb and Trithorax: 70 Years and Counting. Cell 171, (2017).

27. Margueron, R. et al. Ezh1 and Ezh2 Maintain Repressive Chromatin through Different Mechanisms. Molecular Cell 32, 503–518 (2008).

28. de Napoles, M. et al. Polycomb Group Proteins Ring1A/B Link Ubiquitylation of Histone H2A to Heritable Gene Silencing and X Inactivation. Developmental Cell 7, 663–676 (2004).

29. Uhlén, M. et al. Tissue-based map of the human proteome. Science 347, (2015).

30. Lein, E. S. et al. Genome-wide atlas of gene expression in the adult mouse brain. Nature 445, 168–176 (2007).

31. de Bie, P., Zaaroor-Regev, D. & Ciechanover, A. Regulation of the Polycomb protein RING1B ubiquitination by USP7. Biochemical and Biophysical Research Communications 400, 389–395 (2010).

32. Valles, G. J., Bezsonova, I., Woodgate, R. & Ashton, N. W. USP7 Is a Master Regulator of Genome Stability. Front. Cell Dev. Biol. 8, (2020).

33. Cao, Q. et al. The central role of EED in the orchestration of polycomb group complexes. Nat Commun 5, 3127 (2014).

34. López, Y., Nakai, K. & Patil, A. HitPredict version 4: comprehensive reliability scoring of physical protein–protein interactions from more than 100 species. Database (Oxford) 2015, (2015).

35. Lee, A. K. & Potts, P. R. A Comprehensive Guide to the MAGE Family of Ubiquitin Ligases. Journal of Molecular Biology 429, 1114–1142 (2017).

36. Simon, J. A. & Kingston, R. E. Occupying Chromatin: Polycomb Mechanisms for Getting to Genomic Targets, Stopping Transcriptional Traffic, and Staying Put. Molecular Cell 49, 808–824 (2013).

37. Aranda, S., Mas, G. & Croce, L. D. Regulation of gene transcription by Polycomb proteins. Science Advances 1, e1500737 (2015).

38. Yang, J. et al. Maged1 co-interacting with CREB through a hexapeptide repeat domain regulates learning and memory in mice. Mol. Neurobiol. 51, 8–18 (2015).

39. Schmidl, C., Rendeiro, A. F., Sheffield, N. C. & Bock, C. ChIPmentation: fast, robust, low-input ChIP-seq for histones and transcription factors. Nat Methods 12, 963–965 (2015).

40. Martinelli, D. C. et al. Expression of C1ql3 in Discrete Neuronal Populations Controls Efferent Synapse Numbers and Diverse Behaviors. Neuron 91, 1034–1051 (2016).

41. Huggett, S. B. & Stallings, M. C. Genetic Architecture and Molecular Neuropathology of Human Cocaine Addiction. J. Neurosci. 40, 5300–5313 (2020).

42. Barbour, H., Daou, S., Hendzel, M. & Affar, E. B. Polycomb group-mediated histone H2A monoubiquitination in epigenome regulation and nuclear processes. Nature Communications 11, 5947 (2020).

43. Johnson, B. A. et al. Topiramate for the Treatment of Cocaine Addiction: A Randomized Clinical Trial. JAMA Psychiatry 70, 1338–1346 (2013).

44. Shorter, D. & Kosten, T. R. Novel pharmacotherapeutic treatments for cocaine addiction. BMC Med 9, 119 (2011).

45. Desroses, M. & Altun, M. The Next Step Forward in Ubiquitin-Specific Protease 7 Selective Inhibition. Cell Chem Biol 24, 1429–1431 (2017).

46. Lamberto, I. et al. Structure-Guided Development of a Potent and Selective Non-covalent Active-Site Inhibitor of USP7. Cell Chem Biol 24, 1490–1500.e11 (2017).

47. Pozhidaeva, A. et al. USP7-Specific Inhibitors Target and Modify the Enzyme’s Active Site via Distinct Chemical Mechanisms. Cell Chemical Biology 24, 1501–1512.e5 (2017).

## References

48. Bertrand, M. J. M. et al. NRAGE, a p75NTR adaptor protein, is required for developmental apoptosis in vivo. Cell Death Differ. 15, 1921–1929 (2008).

49. Mouri, A. et al. MAGE-D1 regulates expression of depression-like behavior through serotonin transporter ubiquitylation. J. Neurosci. 32, 4562–4580 (2012).

50. Dombret, C. et al. Loss of Maged1 results in obesity, deficits of social interactions, impaired sexual behavior and severe alteration of mature oxytocin production in the hypothalamus. Hum. Mol. Genet. 21, 4703–4717 (2012).

51. Gorski, J. A. et al. Cortical Excitatory Neurons and Glia, But Not GABAergic Neurons, Are Produced in the Emx1-Expressing Lineage. J. Neurosci. 22, 6309–6314 (2002).

52. Haggerty, D. L., Grecco, G. G., Reeves, K. C. & Atwood, B. Adeno-Associated Viral Vectors in Neuroscience Research. Molecular Therapy - Methods & Clinical Development 17, 69–82 (2020).

53. Dassesse, D. et al. Functional striatal hypodopaminergic activity in mice lacking adenosine A2A receptors. J Neurochem 78, 183–198 (2001).

54. Ena, S. L., De Backer, J.-F., Schiffmann, S. N. & de Kerchove d’Exaerde, A. FACS array profiling identifies Ecto-5’ nucleotidase as a striatopallidal neuron-specific gene involved in striatal-dependent learning. J. Neurosci. 33, 8794–8809 (2013).

55. Karadurmus, D. et al. GPRIN3 Controls Neuronal Excitability, Morphology, and Striatal-Dependent Behaviors in the Indirect Pathway of the Striatum. J Neurosci 39, 7513–7528 (2019).

56. Polleux, F. & Ghosh, A. The slice overlay assay: a versatile tool to study the influence of extracellular signals on neuronal development. Sci STKE 2002, pl9 (2002).

57. Dobin, A. et al. STAR: ultrafast universal RNA-seq aligner. Bioinformatics 29, 15–21 (2013).

58. Lacoste, N. et al. Mislocalization of the Centromeric Histone Variant CenH3/CENP-A in Human Cells Depends on the Chaperone DAXX. Molecular Cell 53, 631–644 (2014).

59. Chiva, C. et al. QCloud: A cloud-based quality control system for mass spectrometry-based proteomics laboratories. PLoS One 13, e0189209 (2018).

60. Ran, F. A. et al. Genome engineering using the CRISPR-Cas9 system. Nat Protoc 8, 2281–2308 (2013).

61. Hughes, C. S. et al. Ultrasensitive proteome analysis using paramagnetic bead technology. Mol Syst Biol 10, 757 (2014).

62. Hughes, C. S. et al. Single-pot, solid-phase-enhanced sample preparation for proteomics experiments. Nat Protoc 14, 68–85 (2019).

63. Dayon, L. et al. Relative quantification of proteins in human cerebrospinal fluids by MS/MS using 6-plex isobaric tags. Anal Chem 80, 2921–2931 (2008).

64. Reichel, M. et al. In Planta Determination of the mRNA-Binding Proteome of Arabidopsis Etiolated Seedlings. Plant Cell 28, 2435–2452 (2016).

65. Franken, H. et al. Thermal proteome profiling for unbiased identification of direct and indirect drug targets using multiplexed quantitative mass spectrometry. Nat Protoc 10, 1567–1593 (2015).

66. Ritchie, M. E. et al. limma powers differential expression analyses for RNA-sequencing and microarray studies. Nucleic Acids Res 43, e47 (2015).

67. Huber, W., von Heydebreck, A., Sültmann, H., Poustka, A. & Vingron, M. Variance stabilization applied to microarray data calibration and to the quantification of differential expression. Bioinformatics 18 Suppl 1, S96–104 (2002).

68. Afgan, E. et al. The Galaxy platform for accessible, reproducible and collaborative biomedical analyses: 2018 update. Nucleic Acids Research 46, W537–W544 (2018).

69. Martin, M. Cutadapt removes adapter sequences from high-throughput sequencing reads. EMBnet.journal 17, 10–12 (2011).

70. Langmead, B. & Salzberg, S. L. Fast gapped-read alignment with Bowtie 2. Nat Methods 9, 357–359 (2012).

71. Zhang, Y. et al. Model-based analysis of ChIP-Seq (MACS). Genome Biol 9, R137 (2008).

72. Lawrence, M. et al. Software for Computing and Annotating Genomic Ranges. PLOS Computational Biology 9, e1003118 (2013).

73. Welch, R. P. et al. ChIP-Enrich: gene set enrichment testing for ChIP-seq data. Nucleic Acids Research 42, e105 (2014).

74. Kallin, E. M. et al. Genome-Wide uH2A Localization Analysis Highlights Bmi1-Dependent Deposition of the Mark at Repressed Genes. PLOS Genetics 5, e1000506 (2009).

75. Yang, W. et al. The histone H2A deubiquitinase Usp16 regulates embryonic stem cell gene expression and lineage commitment. Nat Commun 5, 3818 (2014).

76. Zhang, H., Roberts, D. N. & Cairns, B. R. Genome-Wide Dynamics of Htz1, a Histone H2A Variant that Poises Repressed/Basal Promoters for Activation through Histone Loss. Cell 123, 219–231 (2005).

77. Wu, T. et al. clusterProfiler 4.0: A universal enrichment tool for interpreting omics data. Innovation (Camb) 2, 100141 (2021).

78. Kolberg, L., Raudvere, U., Kuzmin, I., Vilo, J. & Peterson, H. gprofiler2 -- an R package for gene list functional enrichment analysis and namespace conversion toolset g:Profiler. F1000Res 9, ELIXIR-709 (2020).

79. Subramanian, A. et al. Gene set enrichment analysis: A knowledge-based approach for interpreting genome-wide expression profiles. PNAS 102, 15545–15550 (2005).

80. First, M., Spitzer, R., Gibbon, M. & Williams, J. Structured Clinical Interview for DSM-IV Axis I Disorders (Internet). (1996).

81. Cubells, J. F. et al. Rating the severity and character of transient cocaine-induced delusions and hallucinations with a new instrument, the Scale for Assessment of Positive Symptoms for Cocaine-Induced Psychosis (SAPS-CIP). Drug Alcohol Depend 80, 23–33 (2005).

82. Chang, C. C. et al. Second-generation PLINK: rising to the challenge of larger and richer datasets. GigaScience 4, (2015).

83. Marees, A. T. et al. A tutorial on conducting genome-wide association studies: Quality control and statistical analysis. Int J Methods Psychiatr Res 27, e1608 (2018).

84. Das, S. et al. Next-generation genotype imputation service and methods. Nat Genet 48, 1284–1287 (2016).

85. Howie, B., Fuchsberger, C., Stephens, M., Marchini, J. & Abecasis, G. R. Fast and accurate genotype imputation in genome-wide association studies through pre-phasing. Nat Genet 44, 955–959 (2012).

86. Butkiewicz, M. & Bush, W. S. In Silico Functional Annotation of Genomic Variation. Curr Protoc Hum Genet 88, 6.15.1–6.15.17 (2016).

87. Rentzsch, P., Witten, D., Cooper, G. M., Shendure, J. & Kircher, M. CADD: predicting the deleteriousness of variants throughout the human genome. Nucleic Acids Res 47, D886–D894 (2019).

88. Huguet, G. et al. Measuring and Estimating the Effect Sizes of Copy Number Variants on General Intelligence in Community-Based Samples. JAMA Psychiatry 75, 447–457 (2018).

89. Wang, K. et al. PennCNV: an integrated hidden Markov model designed for high-resolution copy number variation detection in whole-genome SNP genotyping data. Genome Res 17, 1665–1674 (2007).

90. Colella, S. et al. QuantiSNP: an Objective Bayes Hidden-Markov Model to detect and accurately map copy number variation using SNP genotyping data. Nucleic Acids Res 35, 2013–2025 (2007).

91. Little, J. et al. STrengthening the REporting of Genetic Association Studies (STREGA)— An Extension of the STROBE Statement. PLOS Medicine 6, e1000022 (2009).

